# Vaccination with *Helicobacter pylori* attachment proteins protects against gastric cancer

**DOI:** 10.1101/2023.05.25.542131

**Authors:** Jeanna A. Bugaytsova, Artem Piddubnyi, Iryna Tkachenko, Lena Rakhimova, Johan Olofsson Edlund, Kaisa Thorell, Harold Marcotte, Anders Lundquist, Karin Schön, Nils Lycke, Sebastian Suerbaum, Christian Schulz, Peter Malfertheiner, Lori M. Hansen, Jay V. Solnick, Roman Moskalenko, Lennart Hammarström, Thomas Borén

## Abstract

Most cases of gastric cancer are caused by chronic *Helicobacter pylori* infection, but the lack of early onco-diagnostics and a high risk for antibiotic resistance hampers early intervention through eradication of *H. pylori* infection by antibiotics. We reported on a protective mechanism where *H. pylori* gastric mucosal attachment can be reduced by natural antibodies that block the binding of its attachment protein BabA. Here we show that challenge infection with *H. pylori* induced response of such blocking antibodies in both human volunteers and in rhesus macaques, that mucosal vaccination with BabA protein antigen induced blocking antibodies in rhesus macaques, and that vaccination in a mouse model induced blocking antibodies that reduced gastric mucosal inflammation, preserved the gastric juice acidity, and fully protected the mice from gastric cancer caused by *H. pylori*.

## INTRODUCTION

Chronic life-long infection by *Helicobacter pylori* is the main cause of severe gastric disease, including gastric cancer. Although duodenal ulcer disease is curable by eradication of the infection ^1,2^, antibiotic treatment affords no cure against gastric cancer. Close to one million individuals are diagnosed annually with gastric cancer, all with poor survival rates ^3–5^. Thus, *H. pylori* infection can cause very different disease outcomes and with low probability for accurate long-term diagnostic prognosis. Patients with gastric cancer carry the virulent “triple-positive” *H. pylori*, which is characterized by the presence of the CagA onco-protein, the VacA cyto-toxin, and the blood group antigen binding attachment (BabA) protein ^6,7^, which is an adhesin that binds to the epithelial ABO/Lewis b (Leb) blood group antigens for tight adherence to the gastric mucosa ^8–11^. BabA is highly polymorphic because of its adaptation in binding preferences for the different human ABO blood group phenotypes and their corresponding carbohydrates (glycans) ^10,12,13^. Thus, BabA rapidly mutates and adapts to the gastric pH in different niches in the stomach and to the rising gastric pH during cancer development through the coordinated increase of its acid sensitivity in Leb-binding ^13^. Despite the many amino acid substitutions that occur in BabA, most individuals that carry *H. pylori* nevertheless manage to develop antibodies that block BabA-mediated Leb-binding and hence protect against gastric disease. In contrast, patients with duodenal ulcer disease (DU) can be critically low in such BabA-blocking antibodies (**Bugaytsova et al, ms 1**). The blocking antibodies perform glycan mimicry through competitive binding to the highly conserved amino acid residues that are normally occupied by the Leb glycan in the BabA carbohydrate binding domain (CBD). This mechanism makes the blocking antibodies essentially resistant to antigenic variations in the *H. pylori* BabA antigen. Blocking of BabA-mediated Leb-binding with a resulting reduction in the mucosal adherence of *H. pylori* will over time alleviate the chronic inflammatory pressure. We show that vaccination of mice with a BabA-based vaccine induces broadly blocking antibodies (bbAbs) that reduce gastric mucosal inflammation, preserve the gastric juice acidity, and protect against gastric cancer caused by *H. pylori*. We believe that this new vaccine offers novel treatment options for individuals at risk for gastric disease and cancer.

## RESULTS

### Induction of bbAbs by challenge infection in human volunteers and in rhesus macaques

Low serum titers of bbAbs against Leb-binding constitute a risk factor for duodenal ulcer disease (***Bugaytsova et al, ms 1, Figure 3***), which suggests the possibility of using immune therapy to boost the levels of such blocking antibodies. To test if experimental *H. pylori* infection can induce immune responses in terms of blocking antibodies in humans, we analyzed sera from 29 volunteers who had participated in a previous *H. pylori* vaccination-challenge study and, importantly, had all tested negative for *H. pylori* at the start of the study by routine diagnostic ELISA, ¹³C urea breath tests, *H. pylori* culturing, and histopathological molecular and immunological analysis ^14^. Tests of the series of volunteers’ sera showed that ∼90% (26/29) of the volunteers responded to the experimental infection with the challenge strain *H. pylori* BCM-300 (of African phylogeny (**Figure S1A**) with high titers of bbAbs (**Figure 1A*i***), ^14,15^. The inhibitory activity was assessed as the serum dilution at which half the *H. pylori* binding to Leb was lost, i.e., the 50% Inhibition Titer (IT50). Six volunteers also responded with distinct bbAb activity, i.e., their sera also blocked Leb binding of phylogenetically distant *H. pylori* such as the 17875/Leb strain (**Figure 1A*ii***). Notably, 5 out 6 of these volunteers induced bbAb that reached or superseded the IT50 = ∼30 level that can constitute a risk factor for duodenal ulcer disease (**Figure 1A*ii***) (***Bugaytsova et al, ms 1, Figure 3***). Unexpectedly, however, four individuals demonstrated pre-challenge titers, which might reflect ongoing but undetected low-level *H. pylori* infections as reported by the sensitive 16S RNA sequencing protocols ^16^ (**Figure 1A*i***). These results show (1) that humans can respond to a challenge infection with induction of protective levels of bbAbs, (2) that the bbAbs against the experimental infection are similar to the majority of individuals and patient cohorts world-wide who carry *H. pylori* infections (***Bugaytsova et al, ms 1, Figure 1***), and (3) that the pre-challenge titer results suggest that the IT50 test of a blood sample collected by a finger stick can identify those individuals who are likely to carry non-detectable infections, in this case 14% (4/29) of *H. pylori*-“negative” individuals.

**Figure 1.**
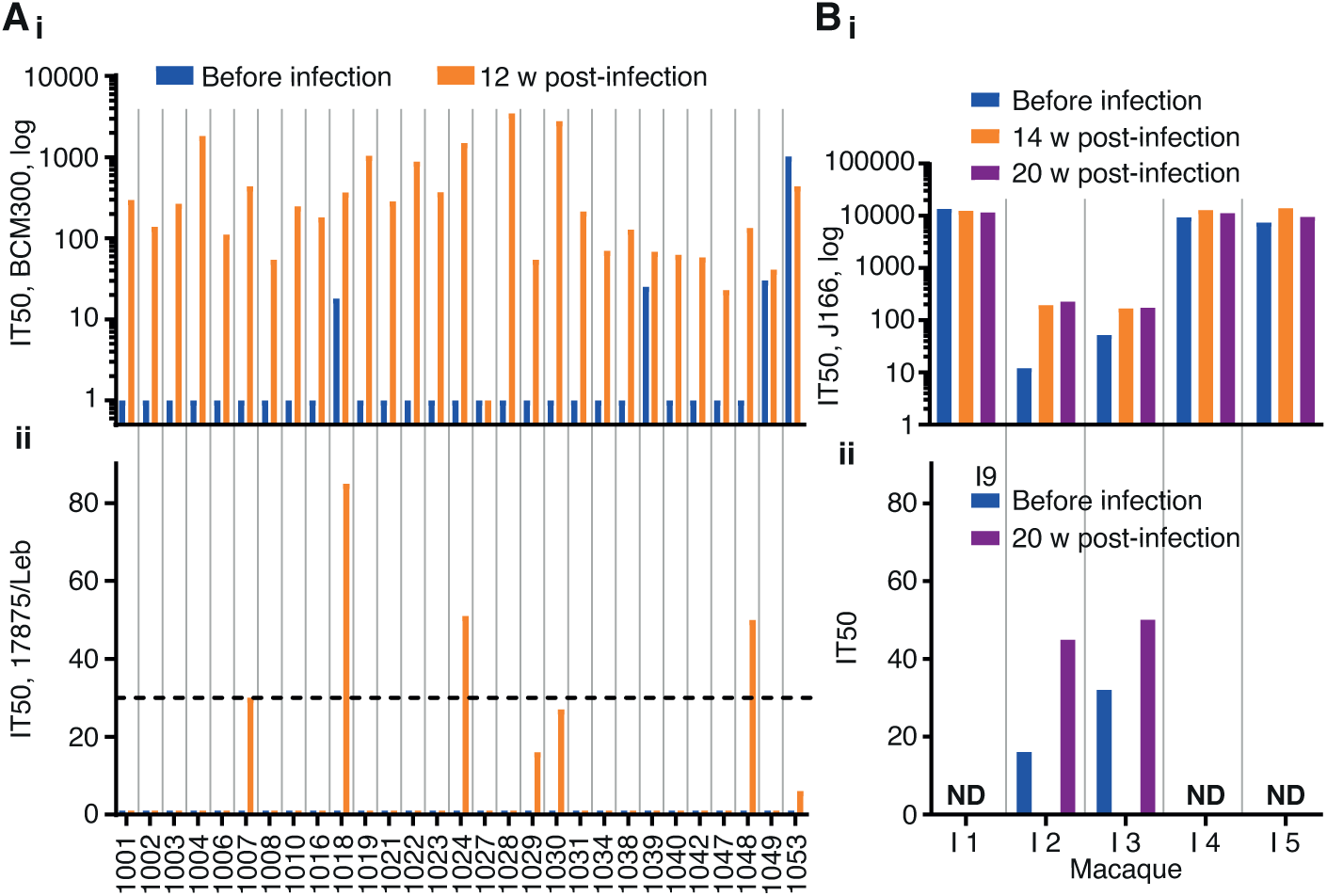
Induction of IT50 responses by challenge infection in humans and rhesus macaques. **(A*i***) The 29 serum samples were tested for IT50s with the challenge strain *H. pylori* BCM300. Only one individual (no. 1027) showed no induction of IT50, whereas four individuals demonstrated pre-challenge titers (blue). (**A*ii***) The unrelated strain 17875/Le identified five serum samples with bbAbs reaching the risk factor IT50 = 30 (dotted line), ranging from an IT50 of 27 to 85, with a mean IT50 of 49 and a median IT50 of 50 (**Table S1A)**. **(B)** Five SPF rhesus macaques were challenge infected with *H. pylori* J166. Two out of the five SPF animals, no. 2 and no. 3, demonstrated induction of an IT50 response, whereas the other three animals demonstrated high pre-challenge IT50 titers (in blue) as tested with J166 (**B*i***) and 17875/Leb (**Figure S1A**) (**B*ii***) Testing of the serum samples from the two animals no. 2 and no. 3 with the phylogenetically distant (unrelated) Indian strain I9 showed that they also induced IT50 responses of bbAbs (**Tables S1B** and **S1C**). ND - Not Determined

The positive results prompted us to test if experimental *H. pylori* infection can similarly induce bbAb responses in rhesus macaques, i.e., in a primate model. From our recent study on BabA adaptation during *H. pylori* infection in rhesus macaques, we analyzed sera from five specific pathogen-free (SPF) animals ^17^. The five animals had all tested negative for *H. pylori* infection, routine diagnostic ELISA, *H. pylori* culturing and histopathology, and they had subsequently been challenge-infected with the *H. pylori* J166 strain ^18^. Sera from 14- and 20-weeks post-infection showed that three of the animals demonstrated high IT50s, whereas two animals exhibited low IT50s, when tested using J166. Unexpectedly, the three animals with high IT50s also demonstrated similarly high pre-challenge titers. These results suggest that these three SPF animals carried non-detectable *H. pylori* infections already at the start of the experiment (**Figure 1B*i***). Notably, the two SPF animals that were clean of pre-titers responded with IT50 = ∼200 when tested with the J166 challenge strain (**Figure 1B*i***) and with a bbAb IT50 = ∼50 when tested with the non-related Indian strain I9 (**Figure 1B*ii***). The two tests demonstrated that experimental *H. pylori* infection can induce similar immune responses with bbAb activity in both humans and rhesus macaques.

### Induction of bbAb responses by vaccination with BabA protein in rhesus macaques

To test if vaccination of primates with BabA as a vaccine antigen can elicit a protective immune response against an *H. pylori* challenge infection, we selected a group of nine SPF rhesus macaques that all tested negative for *H. pylori* infection by routine diagnostic ELISA, *H. pylori* culturing and histopathology. The five animals were immunized intranasally once a week for 4 weeks with a vaccine composed of the native BabA protein purified from *H. pylori* 17875 ^13^ and the mucosal adjuvant CTA1-DD ^19^. The immune response in serum was tested by ELISA at 4 weeks post-vaccination and showed induction of BabA antibodies in 4 of 5 vaccinated animals (**Figure 2A*i***). At the 4 weeks post-immunization time point, all nine animals were challenged with *H. pylori* strain J99, a reference strain of African phylogeny ^20,21^ and hence phylogenetically distant from the 17875 strain of European phylogeny, which was the source of the BabA vaccine antigen (***Bugaytsova, et al, ms1, Figure 2***). The J99 strain infected all animals to similar high H. pylori load as is common in humans and rhesus macaques ^17,22^. Two of the five BabA-vaccinated animals demonstrated reduced infectious loads at 2 weeks after infection, and both “cleared” the infection at 4 weeks. However, at 8 weeks the J99 infection returned although with a two log-fold reduced infectious load as compared to the other three animals (**Figure 2A*ii***). The two animals that responded with a lowered infectious load also demonstrated the strongest ELISA signal (**Figure 2A*i***). These results suggest that the prophylactic vaccination does not protect from infection but that the BabA-ELISA immune response is associated with a temporal reduction of the *H. pylori* infectious load. Next, the serum samples were tested for vaccination-induced IT50 response of bbAb activity. Two animals from the vaccinated group and two animals from the control group demonstrated stable pre-existing IT50s that ranged from 1,000 to almost 100,000, suggesting that they carried a non-detectable *H. pylori* infection already at the start of the experimental series (**Figure S2B**). In contrast, the other animals in each group demonstrated no IT50 pre-vaccination titers. Instead, these vaccinated animals responded with an IT50 = ∼25 at 8 weeks post-challenge by strain J99 and tested by the phylogenetically distant strain 17875 (**Figure 2B**). Notably, testing with the phylogenetically distant Indian strain I9 demonstrated that the animals had a broadly blocking response with an IT50 = ∼100 (**Figure 2B**). The results demonstrate that BabA vaccination can induce bbAb responses with a potential to protect against gastric disease. The many SPF animals with pre-challenge IT50 titers suggests that *H. pylori* can maintain stable long-term BabA expression in rhesus macaques. This might contradict previous reports that *H. pylori* infection in rhesus macaques is accompanied by rapid loss of BabA expression due to loss of the *babA* gene ^23^. However, the two scenarios are principally different because rhesus macaques and humans exhibit similar high infectious loads of *H. pylori* in the range of 10^5^–10^8^ bacteria/gram of gastric tissue. In comparison, the infectious loads in the SPF animals would be in the range of <10^2^/gram i.e., thousands or even a million-fold lower and hence non-detectable with routine diagnostic tools, nor by culture. From this we conclude (1) that measuring IT50s using the ^125^I-Leb-competition technique is a considerably more sensitive method compared to routine *H. pylori* diagnostics; (2) that SPF animals can carry natural *H. pylori* infections; (3) and that the very small infectious loads with no signs of gastric inflammation might support a lifestyle of “humanized” tissue-tropism for *H. pylori* with expression of BabA.

**Figure 2.**
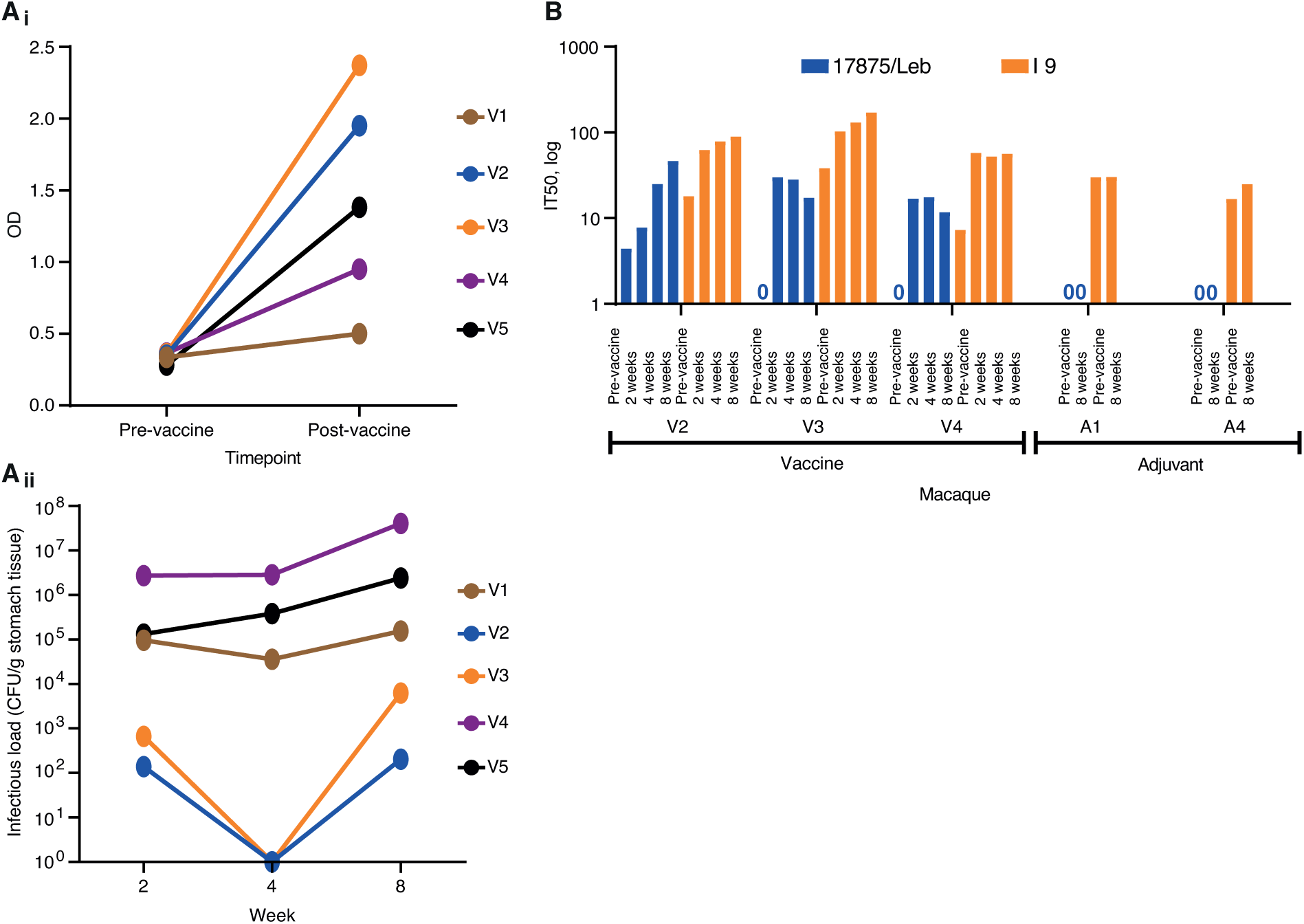
Vaccination and IT50 responses in rhesus macaques. **(A*i*)** ELISA detection of serum BabA antibodies from vaccinated and adjuvant-only control (**Figure S2A*i***) rhesus macaques (**Table S2A)**. **(A*ii*)** Colony forming units (CFUs)/g of stomach-pinch biopsy tissue, i.e., the infectious loads, were tested at 2, 4, and 8 weeks after the infection and were compared to the control animals (**Figure S2A*ii***) (**Table S2B**). (**B)** Test of serum IT50 responses in the vaccinated animals over 2, 4, and 8 weeks with strain 17875 (Europe) and strain I9 (India). The three vaccinated animals no. V2, V3, and V4 responded with *de novo* IT50s when tested with strain 17875. In contrast, animals no. V1, V5, A2, and A3 demonstrated high pre-challenge titers (**Figure S2B**). Notably, the I9 Indian strain demonstrated that the three vaccinated animals no. V2, V3, and V4 also responded with bbAb activity. The I9 strain also confirmed the complete lack of *de novo* IT50s in the non-vaccinated control group (Figure 2B) (**Table S2C**).

### A mouse model of gastric cancer caused by long-term chronic *H. pylori* infection with long-term stability of BabA-mediated Leb-binding activity

There is a recognized epidemiological difference in the development of severe gastric disease where in the majority of individuals the hyper-secretion of acid causes duodenal ulcer disease, whereas a minority of individuals instead initiate the Correa cancer cascade ^24,25^, including the development of atrophy (including loss of the acid-producing parietal cells), stem cell mutations, and resultant gastric cancer ^26^. We recently reported that the *H. pylori* onco-strain USU101 can cause gastric cancer in rhesus macaques ^13,27^. This animal model closely mimics the Correa cascade, but gastric cancer was only found in a single animal after 6 years of *H. pylori* infection ^13^. Thus, we need a model with a faster onset and a higher incidence of severe gastric disease in order to test the efficacy of vaccination for the induction of protective blocking antibodies. Duodenal ulcer disease has not been described in *H. pylori*-infected mice; however, *H. pylori* infected mice that overexpress human gastrin (INS-GAS), which results in excessive gastric acidity, rapidly develop gastric cancer but so do the non-infected INS-GAS mice ^28^. To this end, we developed a mouse model for *H. pylori*-induced gastric cancer. The cancer model is based on the Leb-mouse that expresses a gastric epithelium with humanized Leb-glycosylation, which supports BabA-mediated *H. pylori* attachment ^29^ and gastric mucosal inflammation ^12^ and thus is potentially applicable as a gastric cancer (**Figure 3A**). The Leb-mouse cancer model was developed in a series of steps. First, chronic infection over 12 months by the *H. pylori* onco-strain USU101 resulted in a notably high incidence of gastric cancer of 56% and an additional 18% incidence of dysplasia, i.e., 74% (25/34) of the animals (Group I) developed severe gastric disease with a mean inflammation score of 2.25 (scale 0–3) **(Figure 3B)**. Second, and in sharp contrast, none of the 12 non-infected control Leb-mice (Group III) developed gastric cancer, dysplasia, or inflammation (all scored 0) during the 12-month period (**Figure 3B)**. Thus, gastric cancer and chronic inflammation are entirely dependent on the *H. pylori* infection in the Leb-mouse cancer model. Third, the results were followed up by a second 12-month infection test (Group II), which presented a 40% (12/30) incidence of gastric cancer and an additional 27% (8/30) incidence of dysplasia, i.e., 67% (20/30) of the mice developed malignant gastric disease with a mean inflammation score of 1.4 (**Figure 3B**). Thus, the gastric cancer and dysplasia prevalence was reproducible in the second 12-month test, albeit with a slower transition of the dysplasia into gastric cancer, which is also reflected in the lower mean inflammatory score, i.e., 2.25 vs. 1.4, respectively. All *H. pylori*-infected mice (Groups I and II) demonstrated gastritis at 12 months, which suggests that the take-up of the *H. pylori* infection was ∼100%. At the 12-month end-point, 47% (16/34) of the Group I mice and 30% (9/30) of the Group II mice still carried *H. pylori* infections that were detectable by culture (**Figure 3E**). Unexpectedly, only one out of the 64 infected mice in Groups I and II responded with detectable IT50s during the 12 months of *H. pylori* infection **(Tables S3A** and **S3B**). We conclude that the absence of IT50s of blocking antibodies in mice in response to chronic *H. pylori* infection is very different from the IT50 responses in humans and macaques. The lack of protective IT50s of blocking antibodies in mice against the *H. pylori* onco-strain USU101 seems to be a critical limitation in the Leb-mouse immune system, with resulting excessive chronic mucosal inflammation and an extraordinarily high incidence of gastric cancer and dysplasia.

**Figure 3.**
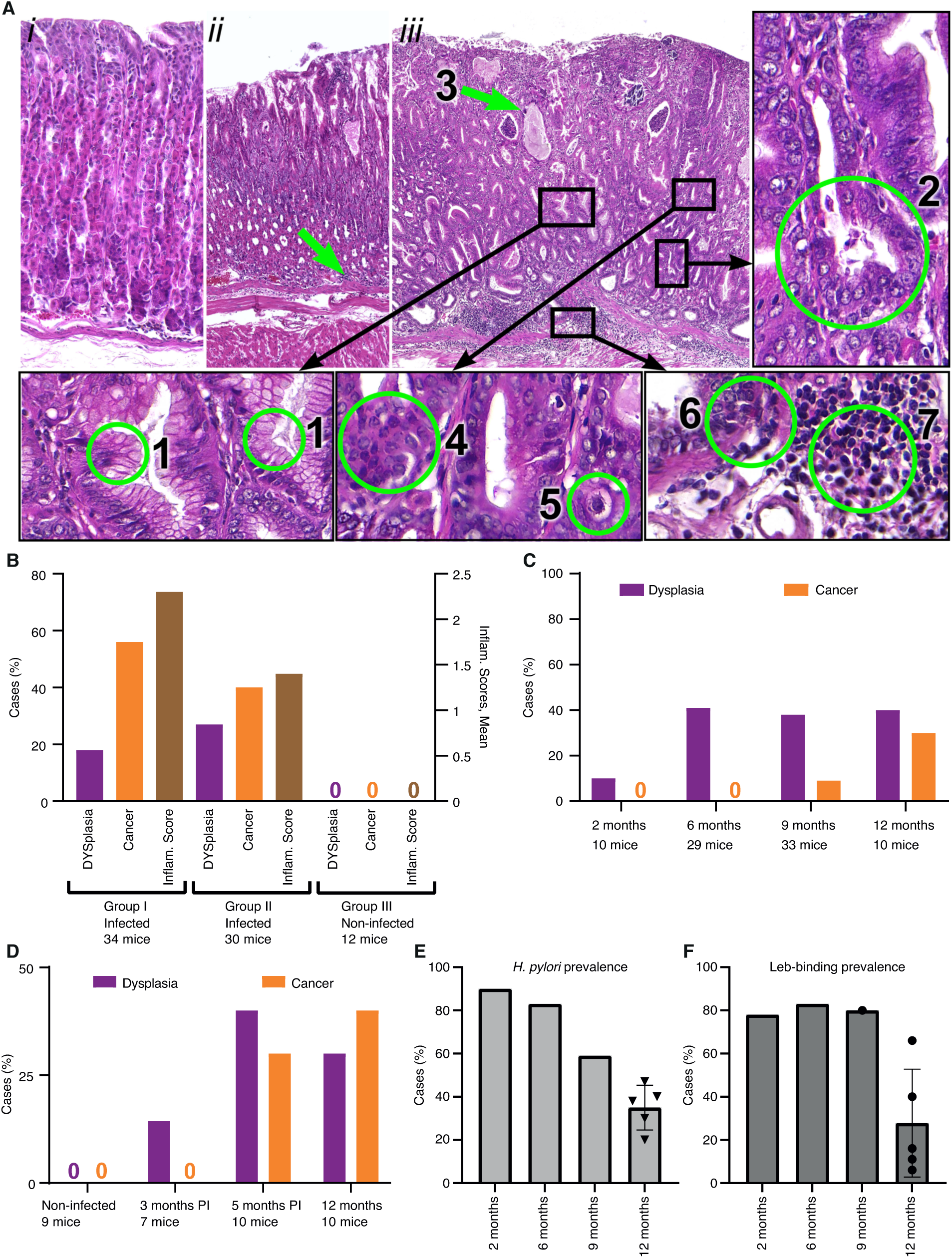
The Leb-mouse gastric cancer animal model. **(A)** The histopathology of the gastric cancer Leb-mouse model was evaluated by H&E-stained sections and blind scoring at the 12-month endpoint according to established criteria ^43^. The mucosal inflammatory infiltration scoring is described in **Figure S3**. (*i*) A non-infected mouse (Group III, no. 2-2, **Table S3C**) with no metaplastic or dysplastic changes and no inflammatory infiltration (scored 0). (*ii*) An infected mouse (Group 1; no. 1-17, **Table S3A**) with no metaplastic or dysplastic changes but with a score of 2 for inflammatory infiltration (green arrow). (*iii*) An infected mouse (Group II; no. 1-5, **Table S3B**) with intestinal metaplasia and numerous goblet cells (1), deformed and branched gastric glands (2), glands with mucus-filled cysts (green arrow, 3), dysplastic glands thickened with layers of nuclei, i.e., cell piling (4), low-grade cellular atypia (5), cancer with growth and penetration through the submucosal layer (6), and cancerous tissue characterized by massive inflammatory cell infiltration (7). Cancer with growth and penetration through the submucosal layer with invasion into the muscular layer is also illustrated in **Figure S3*iii***. **(B)** The Leb-mice were infected with the onco-strain USU101 in two subsequent 12-month periods. Group I included 34 infected mice, Group II included 30 infected mice, and Group III included 12 non-infected control mice. The left Y-axis shows the percentage of mice with dysplasia (lilac) or gastric cancer (orange). The right Y-axis shows the inflammatory infiltration score (brown) **(Tables S3A-C)**. **(C)** The incidence of dysplasia and gastric cancer in the Leb-mouse model after 2, 6, 9, and 12 months of *H. pylori* infection (**Table S3D**). **(D)** The gastric cancer incidence at 12 months in response to antibiotic treatment and eradication of *H. pylori* infection. None of the non-infected mice or antibiotic-treated mice developed cancer at 3 months, and only 1 out of 7 of the treated mice developed dysplasia. In contrast, the mice eradicated of *H. pylori* infection developed similar levels of dysplasia and cancer at 5 months (22 weeks) as the mice that were not treated with antibiotics during the 40-week period (**Table S3E**). **(E)** The *H. pylori* infection was found to be sTable in the mice during the first 6 months of chronic *H. pylori* infection, where the vast majority of Leb-mice were positive for *H. pylori* culture at 2 months (90%) and 6 months (83%) post-infection, with a ∼40–50% reduction at 9 months and 12 months (from (**C**) and **Table S3D**). The 12-month median CFU prevalence of 38% refers to the 12-month infection tests (the non-vaccinated mice) in Figure 3B**, 3C, 5B,** and **5D** (a total of 120 mice, **Table S3F**). **(F)** The BabA-mediated Leb-binding of the *H. pylori* infection was found to be stable in the mice during the first 9 months of chronic *H. pylori* infection, where the vast majority, ∼80%, of the cultured *H. pylori* from (**E**) exhibited preserved Leb-binding capacity (**Table S3D**). The 12-month median and mean Leb-binding prevalence, 16% and 28%, respectively, refers to the series of infection tests of the 120 mice described in (**E**) and **Table S3F**.

#### Identification of the critical age for the Leb-mouse with chronic H. pylori infection to develop dysplasia and/or gastric cancer

To determine the critical age period for establishing gastric cancer in the Leb-mouse model, the animals were scored after 2, 6, 9, and 12 months of chronic infection with the onco-strain USU101. The incidence of dysplasia increased from 6 months after infection, whereas the incidence of gastric cancer increased from 10% at 9 months to 30% at 12 months **(Figure 3C)**. These results show that the development of gastric malignancies in the Leb-mouse model follows the Correa cascade over the life-time of chronic *H. pylori* infection to the final destination, gastric cancer.

#### Identification of the critical age period for the accumulation of the set of mutations that initiate the Correa gastric cancer cascade

We subsequently determined the period in life when the chronic *H. pylori* infection fuels the mucosal inflammatory processes with a critical buildup of mutations that eventually initiates the Correa cancer cascade. For this test, three groups of Leb-mice where infected with the onco-strain USU101. After 12 and 22 weeks the infections were eradicated by antibiotics, whereas the third group was left untreated i.e., with life-long 48 weeks of *H. pylori* infection. The 22-week antibiotic-treated and the 48-week non-treated mice both demonstrated a high (70%) incidence of malignant cell development, i.e., gastric cancer and dysplasia (**Figure 3D).** These results suggest that the 10 weeks infection period, weeks 12 to 22, is critical for accumulation of mutations. These results argue for early intervention in protecting against excessive chronic inflammation because the Correa gastric cancer cascade is the consequence of mutations that occurred earlier in life.

#### Long-term stability of both the chronic H. pylori infection and BabA-mediated Leb-binding activity

Within Group II (**Figure 3B**) and the non-antibiotic-treated group (the 48 weeks infection group) (**Figure 3D)**, only 30% and 20% of the mice carried *H. pylori* at 12 months, respectively. A low prevalence of *H. pylori* infection in patients with gastric cancer is well described ^30^ and is a consequence of the dramatic changes in the gastric environment due to loss of the acid-secreting parietal cells and the resulting rise in gastric pH. To test the stability of *H. pylori* infection during chronic infection, we first analyzed the groups of mice infected with onco-strain USUS101, and after 2, 6 and 9 months (**Figure 3C**) they demonstrated 90% (9/10), 83% (25/30), and 59% (20/34) infection rates, respectively. At the 12 months end-point, the 4^th^ group demonstrated the expected reduction of 40% (4/10) infection prevalence, which was similar to the collective median (38%) of the five different 12-month infections (120 mice) we report here (**Figure 3E**). Thus, we conclude that during the critical period of accumulation of stem-cell mutations, i.e., the first 3–5 months of the experiment, the vast majority of the Leb-mice carry stable *H. pylori* infections. Second, to test for the retained prevalence of BabA-mediated Leb binding activity, we tested the mice that carried *H. pylori* at 2, 6, and 9 months, and ∼80% demonstrated retained BabA-mediated Leb-binding activity (**Figure 3F**). However, at 12 months, i.e., at the Correa cascade end-point, the majority (∼75%) of the *H. pylori* infections in the mice had lost their Leb-binding activity (**Figure 3F**). Thus, ∼70% of the Leb-mice carried *H. pylori* infections that expressed BabA-mediated Leb-binding activity during their first 6 months of life, i.e., the window in time when the critical mutations accumulate that drive the Correa gastric cancer cascade.

### The first vaccine experiment: induction of bbAb response by vaccination and protection against gastric cancer

Our results showed that vaccination with BabA induces IT50s of bbAb activity in rhesus macaques (**Figure 2**). However, for the design of a general vaccine composition we need to take into consideration that long-term *H. pylori* infections in rhesus macaques, mice, and gerbils can reduce the expression of BabA and upregulate the closely related BabB paralog that has an as yet unknown function ^23,31^. In comparison, the strong 84% stability in Leb binding over 6 months of *H. pylori* infection by the onco-strain USU101 in the Leb-mouse (**Figure 3F**) might be a consequence of its “humanized” Leb-glycosylated gastric mucosa that supports BabA-mediated adherence of the *H. pylori* infection. By so doing, the “humanized” Leb gastric mucosa makes the Leb-mouse suitable for vaccine studies. Thus, to test if vaccination can induce protective IT50s, we combined the major *H. pylori* adhesin proteins BabA, BabB ^9^, and SabA ^32^ with the mucosal adjuvant CTA1-DD ^19^ into a vaccine composition. A group of 30 Leb-mice were therapeutically vaccinated one month after established infection by the onco-strain USU101. At the 12-month end-point, both the total incidence of malignant cell development and the inflammation scores were similar in the vaccinated and non-vaccinated (control) mice (77% vs. 67% cancer rates and 1.5 vs. 1.4 inflammation scores, respectively) (**Figure 4A, Tables S3B** and **S4A**). However, the IT50 responses were very differently distributed in the vaccinated group, where the HIGH IT50 mice demonstrated a ∼6-fold reduced incidence of gastric cancer (p < 0.038*) and a ∼9-fold reduction in total dysplasia and cancer (p < 0.033*) (**Figure 4C** and **Table S4B**). The HIGH IT50 mice also scored low (1.0) and were protected from inflammatory infiltration in contrast to the LOW IT50 mice and the non-vaccinated mice, which all scored high for inflammation (LOW IT50, 1.7; Group I, 2.25; and Group II, 1.4) (**Figure 4C** and **3B** and **Tables S4A**, **S3A**, and **S3B**). The high levels of inflammation in the mice with low IT50s is explained by the very strong correlation between high levels of inflammation and gastric cancer (**Figure 4D**), i.e., it is very similar to the high inflammatory infiltration seen in human gastric cancer ^33,34^. Tests by immunoblots showed that only the serum antibodies from the HIGH IT50 mice recognized structural epitopes in BabA (**Figure S4F**), whereas sera from all LOW IT50 and HIGH IT50 vaccinated mice recognized linear epitopes in both BabA and BabB to the same extent, and *vice versa,* where the non-vaccinated animals’ sera showed no BabA/B reactivity at all (**Figures S4A-E**). Next, *H. pylori* was grown from biopsies to understand if the vaccine could provide eradication and sterile clearance of the infection. However, the prevalence of vaccinated mice with LOW IT50 or HIGH IT50 that carried *H. pylori* infection at 12 months were similar in the two groups (9/23 = 39% vs. 2/7 = 28%, and 36% in total) (**Table S4A**).

**Figure 4.**
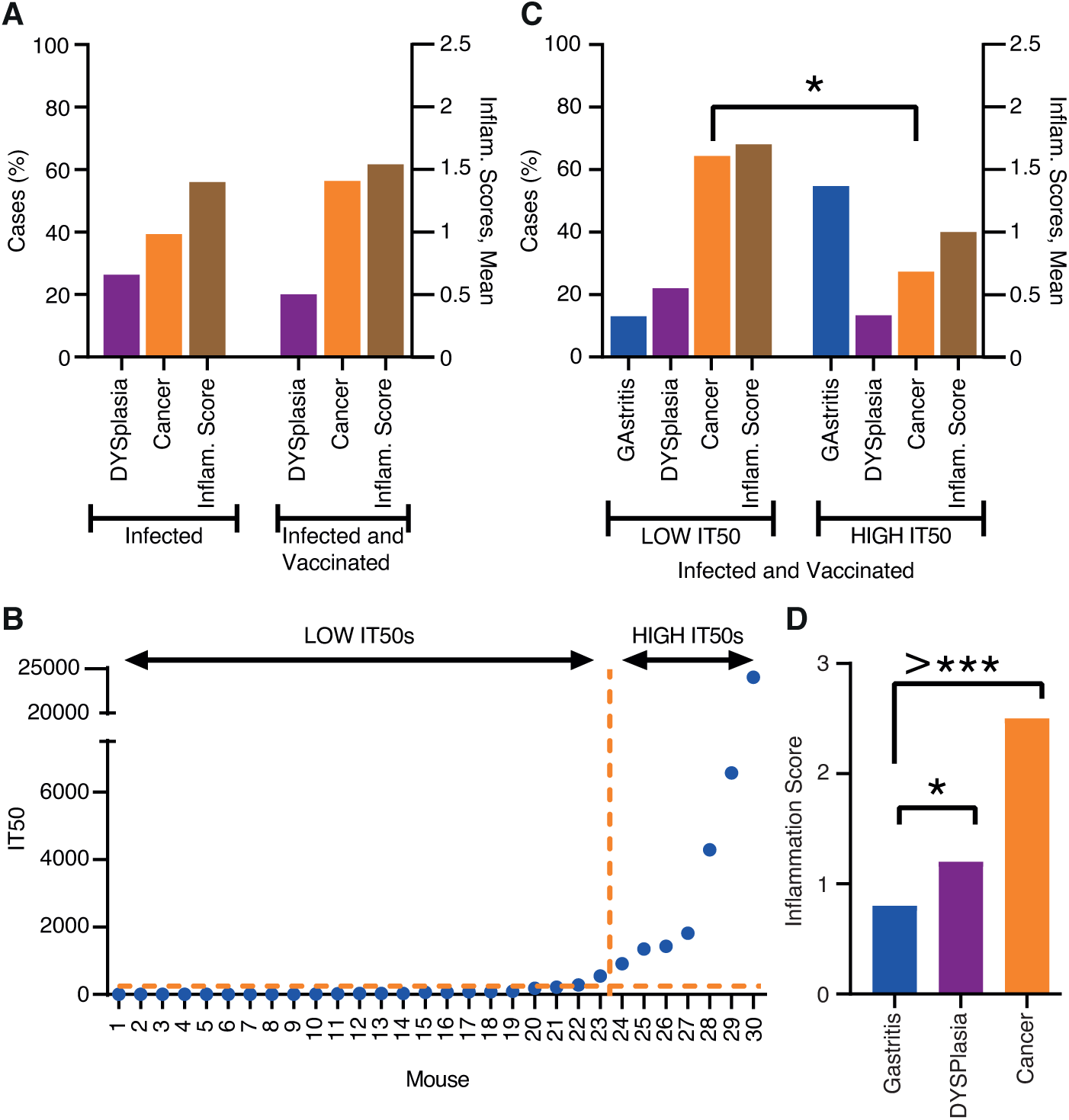
The first vaccination experiment induced protection against inflammation and cancer. **(A)** Two groups, each with 30 Leb-mice, were infected with the onco-strain USU101. One group was therapeutically vaccinated one month later, and all mice were evaluated at the 12-month endpoint. The left Y-axis shows the percentage of mice with dysplasia (lilac) or gastric cancer (orange). The right Y-axis shows the inflammatory infiltration (brown). **(B)** The 30 immunized mice presented in increasing order of IT50, which made a natural divider of the two groups at IT50 = ∼1000 i.e., LOW IT50 vs. HIGH IT50 mice (**Table S4A**). The horizontal hashed line indicates the mean background IT50 = <10 derived from the Group II USU101-infected but non-immunized animals (**Table S3B**). **(C)** Incidence of gastric disease in the mice with LOW IT50 vs. HIGH IT50. The left Y-axis shows the percentage of mice with gastritis (blue) or dysplasia (lilac) or gastric cancer (orange). The right Y-axis shows the inflammatory infiltration (brown) (**Table S4A**). **(D)** The inflammation infiltration score was only 0.8 for gastritis, but increased to 1.2 for dysplasia (p = 0.05*) and almost doubled to 2.5 for the mice with gastric cancer (p = 0.00000000005) (**Table S4C**).

The first therapeutic vaccination pilot experiment suggests (1) that the therapeutic vaccination induces IT50s that reduce gastric mucosal inflammation and gastric cancer, (2) that the non-vaccinated Leb-mice do not respond with protective IT50s during the 12 months of chronic *H. pylori* infection, (3) that the therapeutic vaccination does not cause sterile clearance of the *H. pylori* infection, and (4) that only the protective serum antibodies from the HIGH IT50 mice recognize structural epitopes in BabA.

### The second vaccine experiment: induction of bbAb response by vaccination and protection against gastric cancer

We next performed a second vaccination experiment aimed at increasing the prevalence of HIGH IT50 responders. Thus, mice with an established *H. pylori* infection were therapeutically vaccinated with freshly prepared recombinant BabA, BabB, and SabA antigens. This time, all mice responded to vaccination and with 4-fold higher median IT50 levels compared to the previous HIGH IT50 mouse group (**Figure 5A** and **S5B** and **Table S4D**). Notably, all 21 of the vaccinated mice were protected from gastric cancer, whereas 30% (5/16) of the non-vaccinated mice developed gastric cancer (Fisher: p < 0.01**) (**Figure 5B**). The vaccine also reduced the inflammatory infiltration from 1.5 down to 1.05 (Wilcoxon: p = 0.03*) (**Figure 5C**). Again, the inflammatory infiltration correlated with the severity of disease, and the mice with gastritis had a mean score of 1.1 compared to a score of 2.2 in those with gastric cancer (p < 0.001**). The non-vaccinated mice with dysplasia scored higher (1.67) compared to the gastritis group (Dunn’s test; p = 0.039). In contrast, the vaccinated mice with dysplasia scored only 1.2, which was similar to the score of 1.1 for the gastritis group (**Figure S5A**). The number of vaccinated mice that carried an *H. pylori* infection at 12 months was identical to the non-vaccinated group at 8/21 and 6/16, respectively, i.e., 38% in both groups, and thus essentially identical to the first pilot vaccination experiment (36%) and similar to Group I (47%), Group II (30%), and the 2–12-month test (40%) (**Figure 3E** and **Table S3F)**.

**Figure 5.**
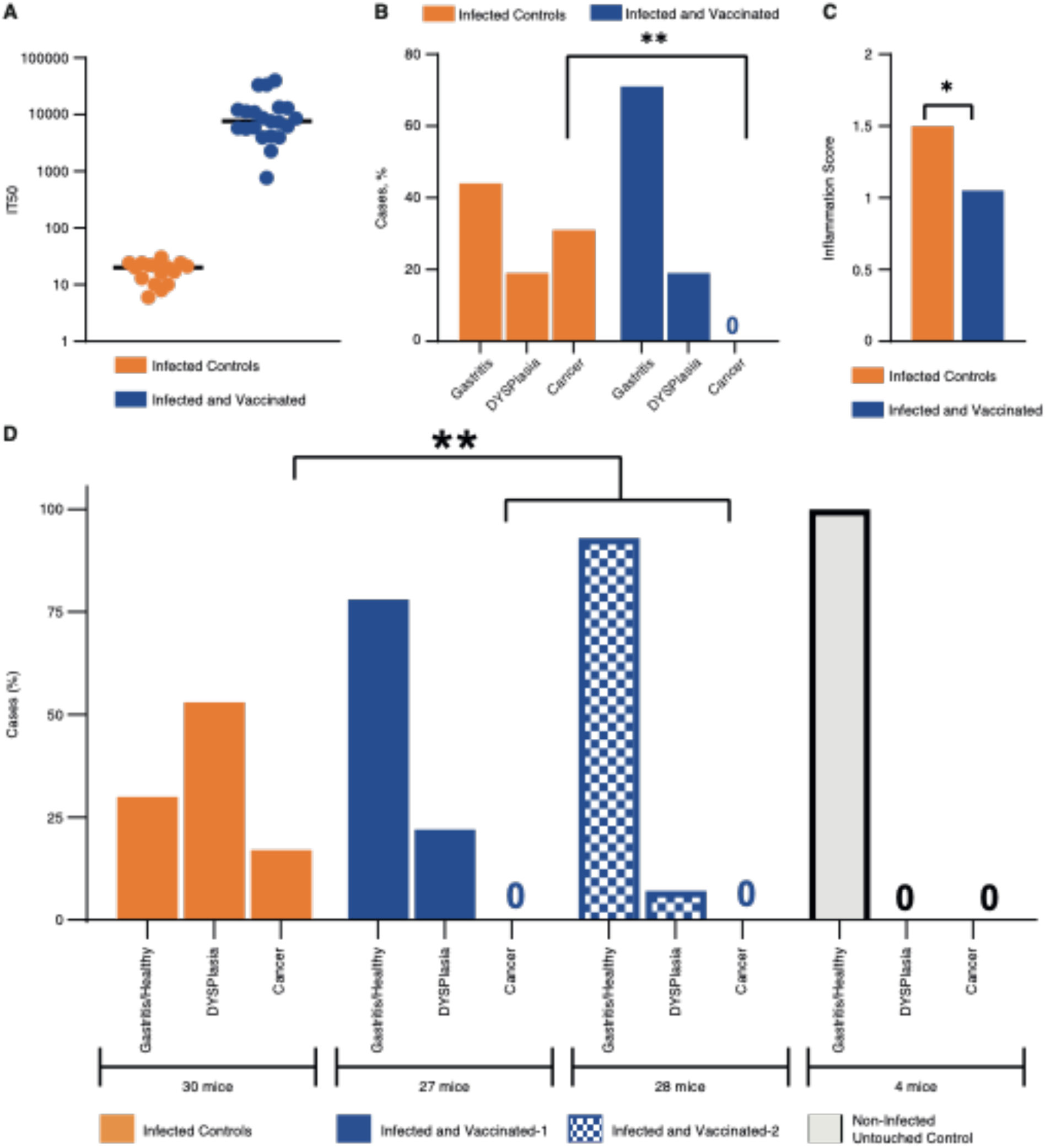
Both the second and third vaccination experiments induced protection against inflammation and cancer. In the second vaccination experiment a group of 37 Leb-mice were infected with the onco-strain USU101, and 21 mice were therapeutically vaccinated one month later and evaluated at the 12-month endpoint. **(A)** The 16 non-vaccinated (orange) control mice demonstrated no IT50 responses compared to the several log-fold higher IT50 responses in the 21 vaccinated (blue) mice, as tested with 17875/Leb. **(B)** The 21 vaccinated mice demonstrated no cancer in contrast to 33% of the non-vaccinated mice (5/16) (p < 0.01**). Four out of the five mice with gastric cancer developed invasive cancer. **(C)** The mean scores for gastric mucosal inflammatory infiltration. The non-vaccinated mice had a higher mean inflammation score of 1.5 vs. 1.05 (p = 0.03*). **(D)** In the third vaccine experiment, the 55 vaccinated mice (blue/hatched) were all protected against gastric cancer, in contrast to the 5/30 (17%) cases of cancer in the non-vaccinated mice (p < 0.0043**). The vaccinated mice exhibited log-fold higher IT50 responses compared to the 30 non-vaccinated mice when tested with 17875/Leb (**Figure S5B** and **Table S5A**).

The second mouse vaccination experiment demonstrated that the therapeutic vaccination can induce high IT50s in 100% of the vaccinated animals, which reduces gastric inflammation and protects against gastric cancer despite not eradicating the *H. pylori* infection.

### The third vaccine experiment: induction of bbAb response by vaccination and protection against gastric cancer

The promising results presented above warranted a combined test of dose-response and reproducibility. Mice were again therapeutically vaccinated, but with High (vaccine-1) or Low (vaccine-2) BabA/BabB antigen levels combined with 5-fold more CTA1-DD adjuvant compared to the previous series. The mice were scored at the 12-month end-point, and the median IT50s were high in both groups and were similar to the second vaccine experiment and were only 2-fold lower in the Low-antigen group (**Figure S5B** and **Table S5A**). The vaccine fully protected against gastric cancer, compared to the 17% (5/30) gastric cancer incidence in the non-vaccinated mice (p < 0.004**). Furthermore, the non-vaccinated group had 57% (16/30) mice with dysplasia, compared to 17% (9/55) of the vaccinated animals (p < 0.0005***) (**Figure 5D**). The High and Low antigen vaccine groups presented 26% (7/27) vs. 7% (2/28) dysplasia, respectively, indicating that a combined high dose of both antigen and adjuvant might be a less protective vaccine composition against malignant development. Notably, the vaccine also protected against total gastric malignant development, with 70% (16 dysplasia + 5 gastric cancer/30) malignancy in the non-vaccinated group compared to only 17% (9 dysplasia/55) in the vaccine groups (p < 0.0001***).

In this third vaccine experiment, the Correa cascade developed somewhat slower with 17% (5/30) gastric cancer cases in the non-vaccinated mice compared to 30% (5/16) in the second vaccine experiment. The slower cascade was also reflected in the proportionally higher incidence of dysplasia of 53% (16/30) compared to the 19% incidence in the second vaccine experiment. The lower gastric cancer incidence might be a consequence of the lower mucosal inflammation level, 1.25, compared to the higher level of 1.5 in the second vaccine experiment test.

The combined cancer reduction in the second and third vaccine experiments, i.e., 5 + 5 cancers in the 46 non-vaccinated mice vs. no gastric cancer cases in the 76 vaccinated mice, indicates that the vaccine is highly protective against cancer (p < 0.001***).

### Vaccination protected against loss of gastric juice acidity

It is well recognized that patients with gastric cancer demonstrate elevated gastric pH due to atrophic gastritis and loss or reduction of the acid-producing parietal cells, which are pathognomonic events in the Correa cascade. To test for similar changes, we measured the gastric pH of the Leb-mice at the 12-month endpoint. We found that the gastric pH was surprisingly low in both the healthy-infected-control and healthy-infected-vaccinated mice, pH 1.4 and 1.7, respectively, and that it increased to pH 2.8 in the mice with gastric cancer (p = 0.033*). The non-vaccinated mice with dysplasia had an increase to pH 2.37, whereas the vaccinated animals were protected against the loss of gastric acidity (p = 0.014*) (**Figure 6A**). Similarly, in the third vaccine experiment the non-vaccinated mice demonstrated elevated gastric pH compared to the vaccinated group (p = 0.0007***) (**Figure S6A**).

**Figure 6.**
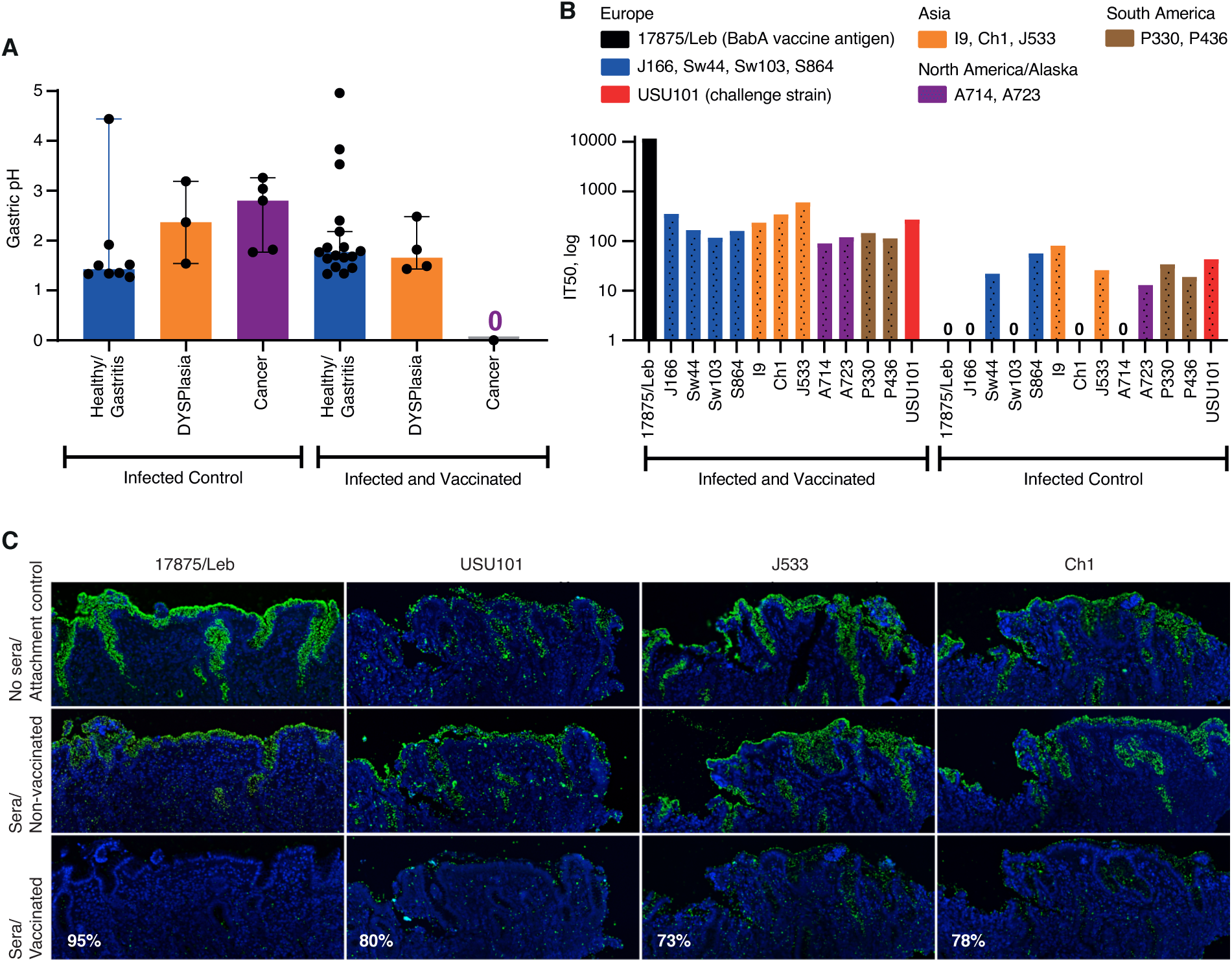
Vaccination preserved gastric acidity, induced bbAb responses, and reduced *H. pylori* gastric mucosal attachment. **(A)**. The non-vaccinated mice with dysplasia or gastric cancer demonstrated elevated gastric pH, whereas the vaccinated group preserved the gastric acidity over the 12-month period (Wilcoxon p = 0.014*) (**Table S4D**). **(B)** The sera of the vaccinated group of mice demonstrated broadly blocking IT50s against the global series of *H. pylori* strains, including the challenge onco-strain USU101. **(C)** The 1:10 diluted sera of the vaccinated group of mice reduced attachment to the human gastric mucosa by (a) 95% for *H. pylori* 17875/Leb, (b) 80% for USU101, (c) 73% for J533 (Japan), and (d) 78% for Ch1 (China) compared to the sera of the non-vaccinated controls (**Figure S6C**).

### Vaccination induced bbAb activity

To test if the vaccination in addition to blocking antibodies also induced responses of protective bbAbs in the Leb-mice, i.e., similar to the bbAb responses in the vaccinated rhesus macaques (**Figure 2**), the sera of the 21 vaccinated mice were tested for IT50s with a full series of 13 global *H. pylori* isolates, including the challenge onco-strain USU101. The serum samples from the vaccinated animals inhibited Leb binding of the full series of *H. pylori* strains in contrast to non-vaccinated controls. The vaccine induced a strong immune response of bbAb activity with a median IT50 = ∼200 (with the antigen source, the 17875/Leb strain, excluded). Thus, the mice responded similar to humans, with generally high IT50 titers of bbAbs against Leb binding with the series of global *H. pylori* infections, including the challenge onco-strain USU101 **(Figure 6B** and **Table S5B**). The results on the general and broadly blocking antibody responses help explain why the BabA vaccine efficiently protects against gastric cancer caused by the phylogenetically distant challenge onco-strain USU101.

### Vaccine induced IT50s of bbAbs against *H. pylori* attachment to the gastric mucosa

The reduced mucosal inflammatory infiltration in the vaccinated animals suggests that the vaccine responses block and reduce *H. pylori* adherence to the gastric mucosa. In support of this notion, the pooled sera of vaccinated mice almost completely (95%) blocked attachment to human gastric mucosa by the strain 17875/Leb. To further understand if the vaccine-induced IT50s also support broadly blocking protection against *H. pylori* attachment to the gastric mucosa, we exposed the challenge onco-strain USU101 to the pooled sera of vaccinated mice. Indeed, the 10-fold dilution of the serum reduced attachment of the mouse model challenge strain USU101 to human gastric mucosa by 80%. The broadly blocking attachment activity of the vaccine was further tested with two *H. pylori* strains from China and Japan, and these notably both demonstrated a similar ∼75% reduction in attachment to the human gastric mucosa **(Figures 6C** and **S6C)**. The efficient broadly blocking attachment of both typical European and Asian *H. pylori* strains to human gastric mucosa suggests that the vaccine holds promise for having global therapeutic and protective efficacy.

## DISCUSSION

A vaccine against *H. pylori* has been the aim although rather a dream during the past 40 years of *H. pylori* research with major efforts by large pharmaceutical and biotech companies and by many research groups. A vaccine is of an even higher priority today due to the increase in multi-resistant *H. pylori* infections that defy current treatment regimens ^35^, and is especially so if the vaccine can reduce the incidence of gastric cancer^36^. Entering the fifth decade of *H. pylori* research, there is still no vaccine on the global market ^37,38^. But why has an efficient *H. pylori* vaccine been so difficult to develop? This is in part because vaccines are by definition prophylactic, and children are commonly vaccinated at an early age before they have encountered the relevant pathogens. However, as we have come to understand, vaccine development against a chronic pathogen such as *H. pylori* is very different because the infection is passed through many generations within the family and thus is inherited by the infants from their parents. Therefore, at whatever age the prophylactic vaccine is provided *H. pylori* is most likely already established in the stomach of the child, although at this early stage the infectious load might be too low for detection using conventional diagnostics tools. Most encouraging, using our new and sensitive IT50 testing method we can identify *H. pylori* by the serum-levels of protective blocking antibodies, both in *H. pylori*-“negative” volunteers and in SPF rhesus macaques. Hence, prophylactic vaccination is less likely to be a useful strategy in protecting against early-age *H. pylori* infections, and our new results instead argue for a delayed therapeutic vaccination, i.e., vaccination of young adult *H. pylori* carriers aimed at attenuating their established *H. pylori* infections.

The second contradiction encountered during the development of *H. pylori* vaccines is that almost every tested antigen in the literature commonly induces log-fold reductions in the infectious load ^39^. However, such log-fold reductions do not deliver sterile clearance of infection because *H. pylori* infects at very high densities of ∼10^5^–10^8^ bacteria/gram of human stomach tissue. Thus, regardless of vaccination, there will be large numbers of *H. pylori* bacteria persisting in the gastric mucosa. In agreement with this, the BabA vaccine could only transiently lower the *H. pylori* infectious load in the rhesus macaque model (**Figure 2**). Thus, our initial vaccine results are in line with the general recognition that human immune responses are not capable of clearing *H. pylori* infections. Fortunately, we can show that mucosal immunization with the BabA vaccine delivers protection against severe gastric disease independently of eradication of the *H. pylori* infection. These results relate to the BabA-dependent adherent lifestyle of *H. pylori* ^13^, which provides the possibilities for therapeutic intervention by vaccine-induced blocking antibodies that competitively reduce *H. pylori* attachment. We suggest that the *H. pylori* pathogen can be turned into a gastric microbiome of more benign and commensal nature by suppression of its inflammatory adherence mode. An attenuated *H. pylori* microbiome might even be beneficial for the individual because childhood eradication of *H. pylori* is correlated with increases in asthma and allergies ^40^ in addition to the reported increased risk for gastrointestinal reflux disease ^41^ and esophageal cancer ^42^.

The new vaccine presented here builds on our results suggesting that bbAbs constitute an Achilles heel for the immune evasion strategy of *H. pylori* infections in humans (**Bugaytsova, et al., ms 1**). Interestingly, in mice the natural immune responses are different because we show that mice do not induce blocking antibodies to chronic *H. pylori* infection, which might be because *H. pylori* is human/primate specific and mice do not carry natural *H. pylori* infections. In contrast, through vaccination the BabA antigen induced responses of bbAbs that also protected the mice from gastric mucosal inflammation and cancer. The efficiency of the vaccine is noteworthy considering the extraordinarily high cancer incidence in the Leb-mouse model, where the majority of mice with chronic *H. pylori* infection develop gastric cancer and/or dysplastic malignant development, i.e., a ∼100-fold higher incidence compared to the clinical situation. **Bugaytsova et al., in ms 1**, showed that the vast majority of individuals that carry *H. pylori* infection already produce bbAbs that perform Leb-glycan mimicry and compete with BabA-mediated Leb-binding. Thus, in humans the vaccine would increase and strengthen the already available natural bbAb levels instead of inducing new and additional immune recognition patterns. In the Leb-mouse gastric cancer model, the vaccine-induced reduction in attachment of the *H. pylori* infection reduces the chronic mucosal inflammation with resulting protection against gastric cancer and dysplasia and preserves gastric juice acidity. Our results suggest translational opportunities for non-antibiotic treatment regimens based on therapeutic vaccination of the group of individuals who demonstrate low and less protective levels of antibodies that will boost their IT50s to reach the critical levels of blocking antibodies needed to provide protection against gastric disease and the silent killer, gastric cancer.

## Abbreviations

bbAb: broadly blocking antibody
CBD: carbohydrate binding domain
GA: gastritis
DYSP: dysplasia
GC: gastric cancer.

## ACKNOWLEDGEMENTS

This paper is in memory of our friend and co-author Dr. Nils Lycke, a great immunologist who contributed importantly to this project. We thank T. Ny for valuable discussions and G. Lisiutin, A. Shevtsova, A. Butsyk, and P. Shubin for excellent technical support; The Biochemical Imaging Centre Umeå (BICU) within the National Microscopy Infrastructure (NMI); The Protein Expertise Platform (PEP) at Umeå University; S. Dübel for providing plasmids; Helicure AB for providing recBabA; and P. G. Falk and J. I. Gordon for the Leb-mouse. We thank M. Borén and E. Morrow for the figure work. This research work was supported by the Swedish Medical Research Council (VR) (2017-02183), Cancerfonden (CF) (CAN 2018/807), the Umeå University Biotechnology Fund, the J.C. Kempe and Seth M. Kempe Memorial Foundation to TB, and the Erling-Persson Family Foundation to TB, LH, and RMo, and by the NIH R21 AI072134 to J.V.S and T.B. The project and was in part performed within The Ukrainian-Swedish Research Center SUMEYA (https://sumeya.med.sumdu.edu.ua/en/). The authors gratefully acknowledge the financial support of the ERASMUS program (EU funding) for PhD student exchange within the collaboration between Umeå University and Sumy State University (grant 2017-1-SE01-KA107-034386).

## AUTHOR CONTRIBUTIONS

All authors contributed substantially to the work and interpretations of the data. J.A.B., A.P., I.T., L.R., J.O.E., K.T., H.M., A.L., L.M.H., J.V.S., and R.M. performed the research and together with L.H. and T.B. analyzed the data. P.M., C.S. and S.S, provided serum samples. N.L. and K.S. provided adjuvant. J.O.E. and A.L. performed statistics. J.A.B., A.P., I.T., J.V.S., R.M., L.H. and T.B. wrote the paper. L.H. and T.B. designed the research. All authors reviewed the manuscript.

## STAR * METHODS

### KEY RESOURCES TABLE

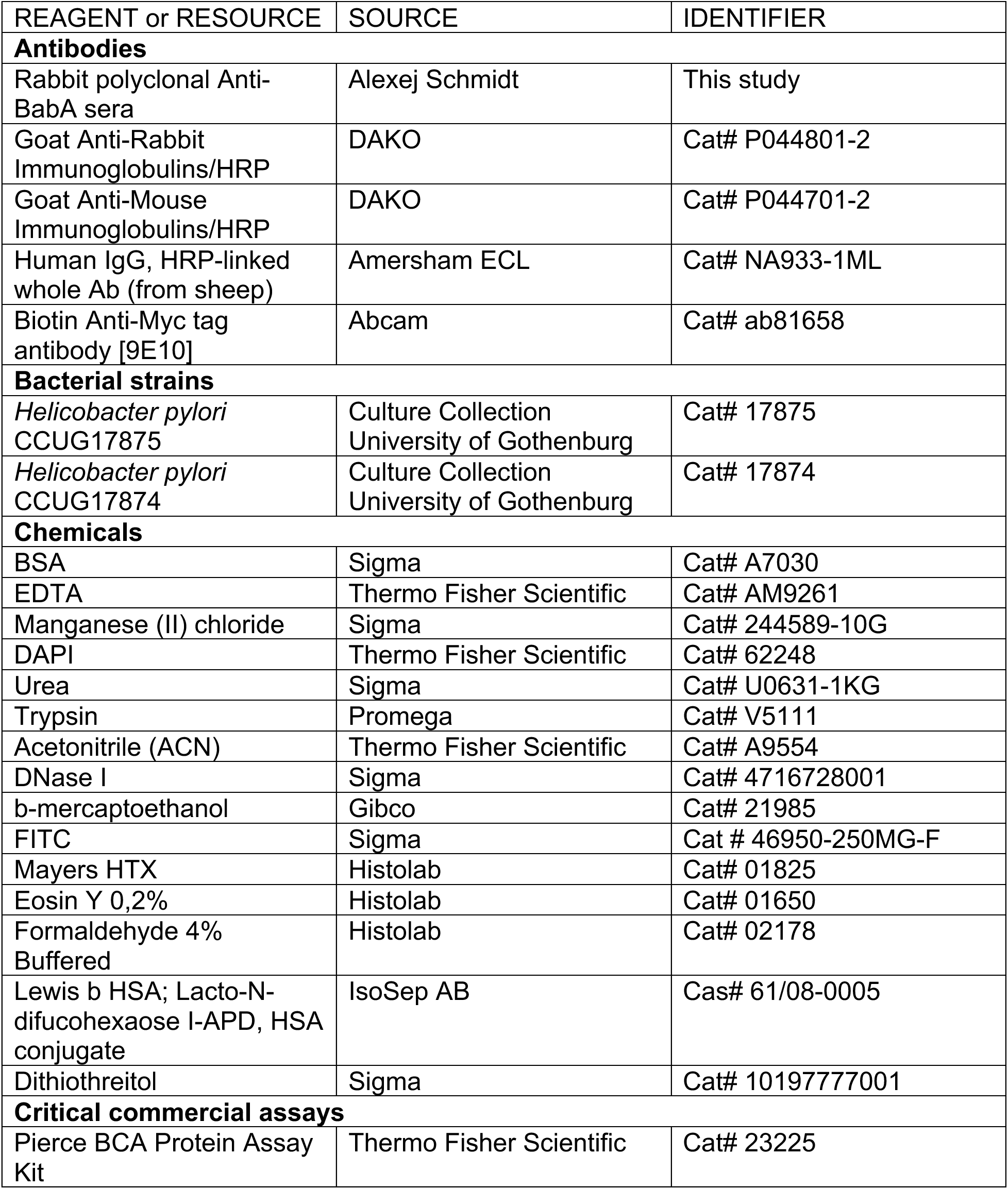
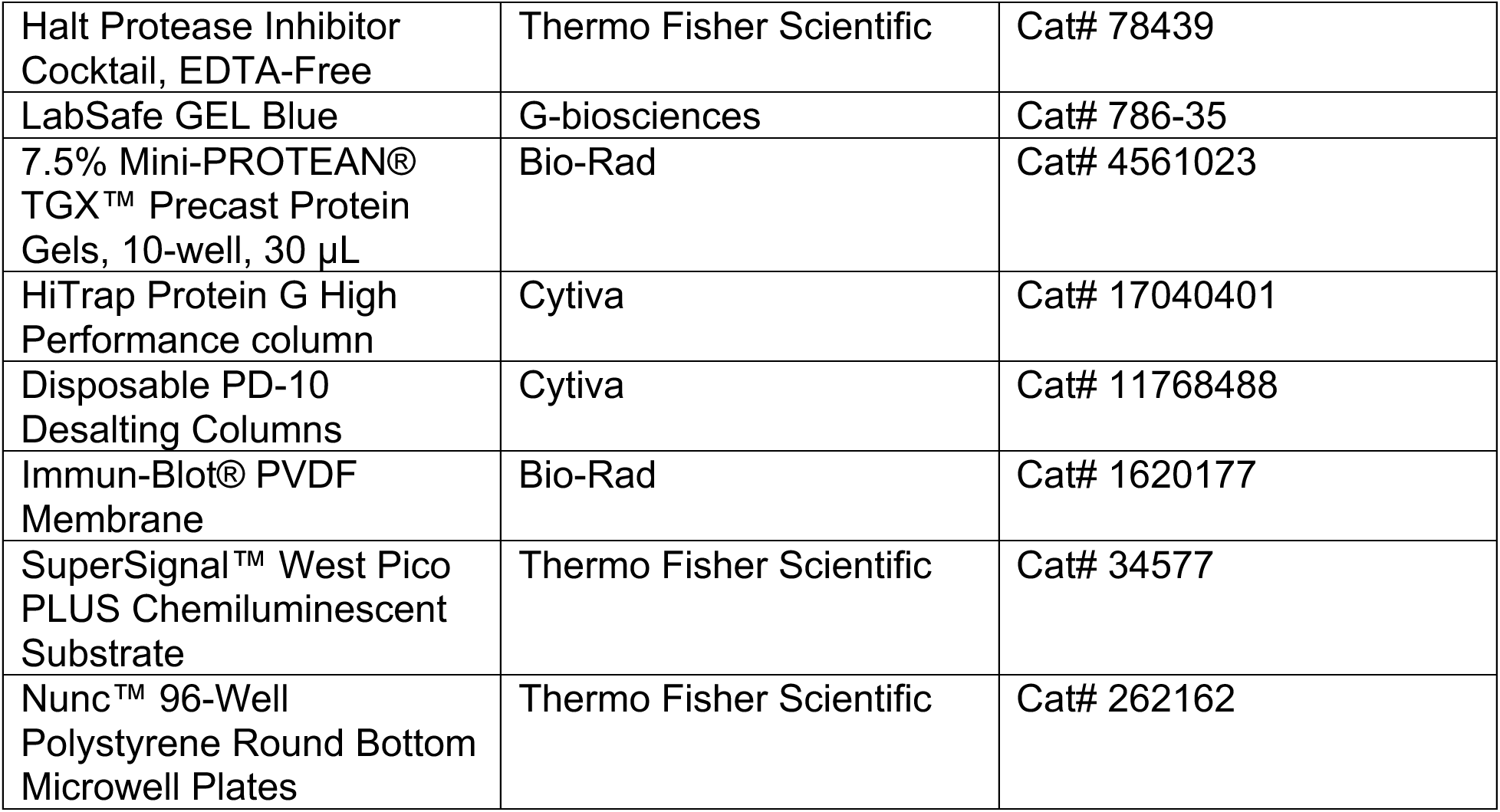

### RESOURCE AVAILABILITY

#### Lead contact

Further information and requests for resources and reagents should be directed to and will be fulfilled by the lead contact, Thomas Borén.

#### Materials availability

Strains and plasmids generated in this study are available upon request to the lead contact, Thomas Borén.

#### Data and code availability

All data are available in the main text or the supplemental information. Any additional information required to reanalyze the data reported in this paper is available from the lead contact upon request.

### EXPERIMENTAL MODEL AND SUBJECT DETAILS

The Supplementary Materials and Methods includes the following topics:

#### 1. *H. pylori* strains

##### A). Laboratory strains

*H. pylori* 17875/Leb is an isolated single clone of *H. pylori* CCUG17875, and it binds to ABO/Leb antigens with high Leb-binding affinity but not to sialylated antigens ^32^. The 17875*babA1A2* strain is a null mutant with the two *babA1* and *babA2* genes deleted from *H. pylori* CCUG17875, referred to as the *babA1A2*-mutant ^9^. *H. pylori* CCUG17874 binds sialylated antigens but not the ABO/Leb antigens ^44^.

##### B). Clinical *H. pylori* isolates

The *H. pylori* isolates Sw44, Sw103, S864, J533, A714, A723, P330, and P436 have been previously described ^10^, and the 13 Indian strains including isolate I9 and the J166 strain have been described in ^13^ and ^18^, respectively.

The *H. pylori* Ch1 strain was isolated from a Chinese individual with a family history of gastric cancer. *H. pylori* BCM-300 ^14,15^ and *H. pylori* USU101 ^13,45^ were as described.

##### C). *H. pylori* culture media

Cultures were performed with blood agar plates, and *H. pylori* cultures were grown in a mixed-gas incubator under micro-aerophilic conditions as described ^44^.

#### 2. Blood group antigen and conjugates

The fucosylated blood group antigen Leb with natural purified oligosaccharides was covalently linked to human serum albumin (HSA) (Isosep AB, Tullinge, Sweden) The Leb-HSA conjugate was used for the radio-immuno assay (RIA) binding experiments.

#### 3. Sera samples

##### 3A. Human sera collection

The Novartis challenge infection study. A total of 29 serum samples were obtained from healthy volunteers who participated in an immunization study with parenteral vaccine against *H. pylori* and challenged with the *H. pylori* BCM-300 strain ^14^. The volunteers all tested negative for *H. pylori* infection by serological, fecal antigen, ^13^C urea breath tests, and gastrointestinal endoscopy with 16 biopsies from the stomach antrum and body for rapid urease testing, *H. pylori* culturing, and histopathological molecular and immunological analysis. Sera were collected before challenge infection and at 12 weeks after (**Table S12**). The series of serum samples from the vaccine clinical study EUDRACT # 2007-003511-31 by Novartis Vaccines and Diagnostics GmbH & Co. KG, Germany, was approved by ethical permit V99P22, 01/08.

##### 3B. Macaque sera, prophylactic vaccination, and challenge infection

###### Challenge infection of the rhesus macaques and tests for sera IT50 (Figure1B)

Animals and the experimental design were essentially according to ^17^. Colony-bred, SPF male and female rhesus macaques between the ages of 2 and 7 years that were free of *H. pylori* infection were derived as previously described ^46^. Four experimental groups were challenged by gavage with 10^9^ CFU/2 ml of *H. pylori* J166 WT (*n*6) as previously described^18^. **IT50 titers** were determined before and at 14 and 20 weeks after infection.

###### Animals and experimental design

Two groups (N = 5 and N = 4) of SPF rhesus macaques were used for the 18-week experiment. The sample size was chosen to detect a 1.8σ difference with 80% power, keeping in mind considerations of animal cost and availability. After identification of SPF macaques by endoscopy (week 0–2), the animals were orally immunized once per week for 4 weeks (week 2–6) with vaccine (CTA1-DD plus BabA) or adjuvant only (CTA1-DD). Four weeks after completion of immunization (week 10), all animals were challenged with rhesus-adapted *H. pylori* J99. Gastric biopsies obtained 2, 4, and 8 weeks after challenge (week 12, 14, and 18, respectively) were used for quantitative *H. pylori* cultures and for histopathology to assess the inflammatory response. Serum and gastric juice were obtained at each endoscopy for determination of BabA-specific antibodies.

###### Immunization

Purified BabA ^13^ (0.50 mg) from *H. pylori* CCUG 17875 and CTA1-DD (0.05 mg) was prepared in 30 mM sodium phosphate buffer (pH 7.0) with 30 mM ocylglucoside. Each animal was given the vaccine weekly for 4 weeks. The vaccine was administered in a total volume of 1.0 ml, half of which was applied slowly to each of the nares with the animal under ketamine (10 mg/g IM) anesthesia.

###### *H. pylori* challenge

All challenges were performed with rhesus-adapted *H. pylori* J99, which has a functional Cag PAI, expresses BabA, and attaches to Leb. Bacteria were grown to early log phase in brucella broth with 5% newborn calf serum in 5% CO2. Animals under ketamine anesthesia (10 mg/kg IM) were oro-gastrically inoculated with 10^9^ cfu.

###### Endoscopy and quantitative cultures

Animals were fasted overnight and given ketamine anesthesia (10 mg/kg IM). The gastric antrum was sampled with four biopsies, which were placed in 200 µl of sterile brucella broth in pre-weighed tubes and homogenized with a sterile glass pestle. An aliquot of 100 µl and dilutions of 10-1 and 10-2 were plated on brucella agar with 5% newborn calf serum to determine the CFU/mg tissue.

###### Determination of BabA antibodies in serum by ELISA

Flat-bottom plates were coated with 500 ng purified BabA per well and blocked with skim milk. Serum was diluted 1:50 in PBS containing 0.05% Tween20 (PBST) and incubated in the wells for 1 h at 37°C. Following washing with PBST, goat anti-monkey IgG-AP was added at a dilution of 1:2000 and incubated as above. Antibodies were detected by the addition of p-nitrophenyl phosphate and measurement of absorbance at 415 nm.

###### Ethics statement

This work was performed at the California National Primate Research Center and the University of California, Davis, in accordance with NIH guidelines, the Animal Welfare Act, and U.S. federal law. All experiments were carried out under protocol 18788 approved by the University of California, Davis, Institutional Animal Care and Use Committee, which is accredited by the Association of Assessment and Accreditation of Laboratory Animal Care. All animals were housed under these guidelines in an accredited research facility fully staffed with trained personnel.

#### 4. Determination of serum Leb-binding inhibition titers (IT50s) (Bugaytsova, ms 1); Blocking buffer and blocking solution

“Blocking buffer” contained 1% BSA (SCA Cohn fraction V) in PBST. For preparation of “SIA-buffer”, 1% BSA was oxidized by 10 mM sodium periodate in 0.1 M acetic acid buffer (pH 4.5) for 1 h. The reaction was stopped by incubating for 30 min in 20 mM sodium bisulfite and 0.125 M potassium phosphate. The solution was then dialyzed against deionized water overnight at 4°C. For SIA blocking buffer, the periodate-treated BSA was mixed with PBS and then filtered through 0.22 µm filters, aliquoted, and kept at –20°C.

##### A). Labeling of Leb-HSA by ^125^I

The Leb-HSA conjugates (IsoSep AB, Tullinge, Sweden) were I^125^-labeled (I^125^-Leb-conjugate) using the chloramine-T method ^44^.

##### Analysis of BabA binding properties by RIA

^125^I-labeled Leb-HSA conjugate (“hot conjugate”) or I^125^-labeled Leb-HSA diluted with unlabeled conjugate (“cocktail”) ^44^ was mixed with 1 mL of bacterial suspension (OD_600_ = 0.1) in blocking buffer. Following incubation, the bacteria were pelleted by centrifugation at 13,000 × *g*, and the ^125^I in the pellet and in the supernatant was measured using a 2470 Wizard^2^ Automatic Gamma counter (PerkinElmer, Waltham, MA, USA) giving a measure of binding activity (% binding).

##### Analysis of sera IT50s by RIA

Serum samples from mice, macaques, and humans were tested for their ability to inhibit the binding of radiolabeled Leb-HSA conjugate (I^125^-Leb-conjugate) to selected *H. pylori* strains. In order to compare IT50s fairly between strains with varying maximum binding properties, all strains were calibrated in a pilot experiment to find the dilution corresponding to 10% I^125^-Leb-conjugate binding. To ensure consistency of bacterial numbers and to aid pellet recovery, strains were diluted with *H. pylori* 17874, which does not bind Leb (carrier strain). For example, 17875/Leb at an OD600nm = 1.0 (2.5 × 10^9^ CFU/mL) was diluted 1:900 with *H. pylori* 17874 to reach 10% binding. Serial dilutions of the serum were made in 60 µL blocking buffer. Bacterial mixtures containing the target strain and carrier strain *H. pylori* 17874 were prepared, and before adding it to the final volume I^125^-Leb conjugate was added to a final concentration of 1 ng/60 µL. A total of 60 µL of the bacterial mixture was then added to a final volume of 120 µL. After addition, the tubes were rotated for 17 h at room temperature. Samples were centrifuged (13,000 × *g* for 13 min), and the I^125^-Leb-conjugates in the pellet and supernatant were measured to determine the amounts of bound and free conjugate, respectively. The relative titer of the tested serum was defined as the dilution titer sufficient to reduce Leb binding to half the maximum value as determined by binding of the Leb conjugate in the absence of serum (IT50).

#### 5. The Leb-mouse

##### Maintenance of FVB/N-Leb mice

*FVB/N* transgenic mice that express human

α-1,3/4-fucosyltransferase and thus have Leb-glycosylated gastric epithelium (Leb-mice) ^29,47^ were used for this study. Th Leb-mice were kept in an IVC (individual ventilated cage) with no more than four animals per cage at the Umeå Center for Comparative Biology (UCCB), Umeå University. The mice were housed in a 12-h dark/light cycle environment with *ad libitum* access to commercial diet formula and tap water.

##### Ethics statement

The mice were maintained by trained personnel at the animal facility of the UCCB under pathogen-free conditions. The Leb-mouse model was approved by ethical permit Dno. A10-2018, A19-18 by the Umeå University ethics committee and complied with the regulations and rules of the Swedish Animal Welfare Agency and with the European Communities’ Council Directive of 22.09.2010 (2010/63/EU).

##### Genotyping of Leb-mouse breeder pairs

The KAPA2G Fast genotyping Mix (KAPA Mouse genotyping kit, Roche Diagnostics) was used for DNA extraction from mouse tissue samples followed by extraction lysis at 75°C for 10 min and 95°C for 5 min. The DNA extracts were diluted 10-fold with 10 mM Tris-HCl (pH 8.0–8.5). PCR: diluted extracts were mixed with PCR master mix (2XKAPA Genotyping Mix with dye), 10 µM Forward primer and 10 µM Reverse primer for human glucosyltransferase ^29^ and Actin (of mouse origin). Cycling protocol for PCR: initial denaturation at 95°C for 3 min, 40 cycles of denaturation at 95°C for 15 sec, annealing at 60°C for 15 sec, and extension at 72°C for 15 sec/kb, and a final extension at 72°C for 1 min. The PCR extracts were separated by 2% agarose gel electrophoresis. All breeder pairs were tested positive for Leb-transgenicity by PCR.

##### Phenotyping of experimental Leb-mice

The tissue sections of mouse gastric mucosa were deparaffinized by submerging into two portions of xylene for 5 min each and rehydrated with a descending gradient of ethanol of 99%-96%-70% for 2 min in each (AnalaR NORMAPUR, Avantor, UK). Antigen retrieval was performed in a 10 mM citrate buffer at pH 6.0 in a 2100 Antigen Retriever (Aptum Biologics Ltd., UK). Samples were blocked with 10% normal goat serum (Thermo Fisher Scientific, USA) for 10 min at room temperature, probed with mouse monoclonal anti-Leb antibodies (Immucor Inc. CA, USA) for 1 h at room temperature followed by washing in PBS-T and developing with goat-anti-mouse Alexa568-conjugated IgG (Thermo Fisher Scientific, USA) for 1 h at room temperature. Nuclei were counterstained with DAPI (Sigma-Aldrich, USA) for 5 min. Each staining procedure included positive controls (PCR-positive mouse gastric tissue with previously estimated expression of Leb in the gastric epithelium) and negative controls (PCR-negative mouse gastric tissue with no Leb expression). The blinded evaluation of the immunostaining was performed by a pathologist on a Leica Thunder microscope (Leica Microsystems, Germany).

##### Mouse surgery

All male mice underwent surgical castration at 4–6 weeks of age due to aggressive behavior in the groups. All manipulations were performed under pathogen-free conditions. Surgical castration of male mice was related to aseptic survival surgery following the principles in ^48^ and the Guidelines for Survival Rodent Surgery (NIH ARAC, 2005, ^49,50^. Prior to surgical invasion, the mouse was anesthetized through the inhalation of 2% Isofluran. Analgesia was provided by Buprenorphine (0.1 mg/kg SC). Fully anesthetized mice were placed in a supine position. After aseptically preparing the surgical field (the abdomen), a vertical incision was made ∼1 cm in the lower abdomen, anterior to the penis, passing through the skin and peritoneum, and the testes were extirpated alternately by a cautery pen. The peritoneum was closed with absorbable sutures and the skin was stapled with surgical wound clips (Reflex Autoclip System, 7 mm). After surgery, the mice were kept individually in clean cages until full recovery (up to 10–14 days). After recovery, the metal clips were removed and the mice were housed in groups of four mice per cage.

##### Infection of Leb-mice

Mice were infected at the age of 8–15 weeks. Mice were infected a total of four times for two consecutive weeks by *per os* gavage with 200 µL of a 50:50 mixture of *H. pylori* strain USU101 and a 12-month mouse-adapted USU101 output. The infecting inoculum was adjusted by Brucella broth to an approximate dose of 10^9^ CFU/mL (OD_600_ = 1.0).

##### *H. pylori* culture for CFUs/g gastric tissue

Stomach samples were transferred into transportation media immediately after sacrifice. Back in the lab, they were vortexed and cultured on selective plates (Brucella agar (BD BBL) – 43 g/L supplemented with Iso-Vitox 1% (v/v), 10% citrated bovine blood (Svenska Labfab), 100 µg/ml vancomycin, 20 µg/ml amphotericin B, 10.7 µg/ml nalidixic acid, 200 µg/ml bacitracin, and 3.3 µg/ml polymyxin B). Plates were incubated under microaerophilic conditions at 37°C for 10 days and examined for bacterial growth. The number of single colonies on the plate was recalculated as CFU/g of gastric stomach tissue sample used for the culture.

#### 6. The Leb-mouse gastric cancer model, part 1

##### Sacrifice of the mice and collection of material

The mice were fasted for 2 h prior to sacrifice by cervical dislocation, and blood samples were taken by heart puncture and kept on ice. Serum samples were separated as soon as the samples arrived at the lab by centrifugation for 10 minutes at 11,500 × *g* at room temperature and were stored at – 80°C until use. The mouse autopsy started from the first cut to remove the ventral skin. A second cut across the midline opened the abdominal cavity, and two subsequent cuts removed the sternum. Terminal blood was collected by cardiac puncture. The stomach was extracted from the abdomen by cutting off 2–3 mm from the esophagus to the gastroduodenal junction. Before opening the stomach, the pH values were measured. The stomach was then opened along the long curvature, emptied of its contents, and weighed. The stomach was divided into two parts, including the forestomach, corpus, and antrum. One part was fixed in 4% PFA (HistoLab, Sweden) for histological examination of the tissue. The mucous layer of second part of the stomach was scraped off with a sterile blade and placed into transport medium (2 g casamino acid (Difco), 2 g peptone (VWR), 0.4 g yeast extract (Merck), 0.32 g bacteriological agar (Acumedia), 0.04 g L-cysteine (Merck), 0.2 g glucose, 28 ml glycerol, 1 g sodium chloride (Merck), and 240 mL Milli-Q (Millipore) filtered water (pH 7.0) for quantitative culture.

##### Gastric pH measurement

The stomach pH was measured in mice fasted for at least 2 h prior to sacrifice using a pH meter (Mettler Toledo Five Easy FE20) with a micro electrode (InLab Micro, Mettler Toledo). Before use, the pH meter was calibrated in the range of pH 2.0 to pH 4.0. To obtain the gastric pH value, the pH electrode was inserted through the pyloric opening in the lumen of stomach without touching the gastric walls. The luminal pH was read when the electrode reached pH stability.

##### Mouse blood collection

After restraining the mouse, blood was collected by submandibular venous puncture with a lancet (Goldenrod animal lancet, 4 mm) ^51^. The blood volume never exceeded 0.2 ml.

##### Mouse serum preparation

The collected whole blood sample was allowed to clot at room temperature for 30 min, and the clot was removed by centrifugation for 11 minutes at 11,500 × *g*. The supernatant, i.e., the serum, was collected in sterile Eppendorf tubes, kept on ice until delivery to the laboratory, and then transferred to a –80°C freezer.

##### Histological analysis of Leb-mouse gastric mucosa

Mice were sacrificed by cervical dislocation and their stomachs were dissected through the small curvature. A representative part of the organ with all anatomical regions (forestomach, corpus, and antrum) was placed in a standard histological cassette (Thermo Fisher Scientific, USA) between two biopsy pads (Thermo Fisher Scientific, USA) in order to prevent tissue deformation. Tissue samples were fixed in a 4% neutral paraformaldehyde aqueous solution (HistoLab, Sweden) for 24 h and saturated with paraffin in a Leica ASP300S tissue processor (Leica Microsystems, Germany). Standard paraffin blocks were made with a Leica EG1140 embedding station or a Leica Arkadia H/C system (Leica Microsystems, Germany) with Histowax paraffin (HistoLab, Sweden). Sections with a thickness of 4 µm were cut with a Leica RM2255 automated microtome (Leica Microsystems, Germany), placed on SuperFrost Plus adhesive slides (Thermo Fisher Scientific, USA), and dried overnight at 37°C. Samples were submerged in two portions of xylene for 5 min each for deparaffinization. Rehydration was performed in absolute ethanol (10 min), 96% ethanol (2 min), and 70% ethanol (2 min). Slides were stained with Mayer’s hematoxylin <u>and</u> eosin (HistoLab, Sweden), mounted with Pertex mounting medium (HistoLab, Sweden), and dried overnight in a fume hood. All slides were scanned with a Pannoramic 250 Flash II scanner (3DHistech, Hungary).

##### Pathological evaluation of gastric disease and inflammation

The hematoxylin and eosin-stained slides contained all anatomical regions of the mouse stomach (forestomach, corpus, and pylorus) and duodenum. Microslides were digitized to full-slide scans with a 250 Flash III tissue scanner (3DHistech, Hungary). All samples were analyzed in the 3DHistech SlideViewer (3DHistech, Hungary). Briefly, at least six fields with an area of 1 mm^2^ of each sample were used for the evaluation. The inflammatory cells (lymphocytes, neutrophils, and macrophages) were counted manually in each field. The mean number of inflammatory cells per field was then graded according to the scale in **Figure S3**. All samples were scored for gastritis, dysplasia, and cancer in situ by a pathologist (R. Mo) in a blinded manner ^13,43^.

#### The Leb-mouse gastric cancer model, part 2

##### Identification of the critical age for the Leb-mouse with chronic *H. pylori* infection to develop dysplasia and/or gastric cancer

###### Experimental design

We sacrificed *H. pylori*-infected mice (60 days old) at different time points to evaluate the gastric cancer incidence and persistence of infection by analysis of stomach biopsies. Mice were sacrificed at 2 months post-infection (n = 10), 6 months post-infection (n = 30), 9 months post-infection (n = 34), and 12 months post-infection (n = 10).

#### The Leb-mouse gastric cancer model, part 3

##### Identification of the critical age period for the accumulation of the set of mutations that initiate the Correa gastric cancer cascade

###### Animal model

For this study, we used 36 FVB/N Leb transgenic male mice that had been surgically castrated at 6–7 weeks of age.

###### Experimental design

A total of 36 mice were distributed randomly into 4 experimental groups: group 1 – infected and treated at 12 weeks post infection, group 2 - infected and treated at 22 weeks post infection, group 3 – infected and not treated, and group 4 – not infected and not treated. All mice were sacrificed at 48 weeks post infection.

###### Treatment

Mice were treated *per os* gavage with a mixture of metronidazole, clarithromycin, and omeprazole for 7 days at 12- or 22-weeks post infection according to the experimental set up. No repeated therapy was provided during the whole period of this study. One dose (200 µL) was given per mouse per day consisting of omeprazole (400 µmol/kg, Sigma Aldrich Lot# LRAC0716), metronidazole (14.2 mg/kg, Sigma Aldrich Lot# SLBQ4358V), and clarithromycin (7.15 mg/kg, Sigma Aldrich Lot# 019M4018V).

#### 7. Vaccination experiments

##### Immunization of Leb-mice with BabA, BabB, and SabA antigens (Figure 3B)

Recombinant proteins were expressed and purified as described ^52^. The vaccination cocktail contained 4 µg (High) or 1µg (Low) of each recombinantly produced antigen (BabA, BabB, and SabA) and 1 or 5 µg of CTA1-DD (cholera toxin-based adjuvant).

Nasal immunization was performed under 4% isoflurane inhalation anesthesia with 20 µl of the antigen cocktail (10 µl in each nostril). The vaccination experiment started on week 6 and was performed once a week in two rounds – the first round was at weeks 7–10 after the beginning of the experiment, and the second round was at weeks 17–20 after the beginning of the experiment.

For the first vaccine experiment, 60 FVB/N Leb-mice infected with *H. pylori* USU 101 were divided into two groups – 1) the infected but not vaccinated controls (n = 30) and 2) the infected and vaccinated test group (n = 30). Mice were immunized with the high dose, 4 µg, of the vaccination cocktail.

For the second vaccine experiment, 37 FVB/N Leb-mice infected with *H. pylori* USU 101 were divided into two groups – 1) the infected but not vaccinated controls (n = 16) and 2) the infected and vaccinated test group (n = 21). Mice were immunized with the high dose, 4 µg, of the vaccination cocktail.

For the third vaccine experiment, 89 FVB/N Leb-mice infected with *H. pylori* USU 101 onco-strain were divided into the following groups: 1) the infected but not vaccinated controls (n = 30), 2) the infected and vaccinated test group with the high dose vaccination cocktail (n = 27), 3) the infected and vaccinated test group with the low dose vaccination cocktail (n = 28), and 4) the non-infected, non-vaccinated (untouched) controls (n = 4).

#### 8. SDS-PAGE and immunoblot detection

##### Polyclonal rabbit sera for immunoblot detection of BabA and BabB

BabA and BabA proteins were detected with the polyclonal anti-BabA VITE antibody (1:6000 dilution) and polyclonal anti-BabB VIRA antibody (1:3000 dilution) and secondary HRP-goat-anti-rabbit antibody diluted 1:1000 (DakoCytomation, Denmark A/S) according to ^13^.

##### Immunoblot detection of denatured BabA (Figures S4A, B, C, D, E) or semi-native BabA (Figure S4F)

SDS-PAGE was performed with 7.5% or 10% Mini-PROTEAN TGX Gels (Bio-Rad Laboratories, Hercules, CA, USA). BabA protein was mixed with Laemmli buffer under non-reducing, semi-native conditions and mild heating at 37°C for 30 min and supplemented with BSA (SCA Cohn fraction V, SWAB, Sweden). For reduced and denaturing conditions, sample buffer was supplemented with reducing agent (5% β-mercaptoethanol) and boiled for 5 min prior to loading onto the gels. For immunoblots, proteins were transferred to a polyvinylidene difluoride membrane (Bio-Rad Laboratories, USA) and blocked with 5% skim milk in TBS-T. Detection of recombinant BabA with sera from immunized mice on immunoblots was as described above, with a serum dilution 1:250 for reduced and denatured protein or a 1:500 dilution for semi-native probe followed by secondary goat-anti-mouse HRP-conjugated goat anti-mouse antibody diluted 1:1000 (DAKO). Signals were developed with ECL chemiluminescence (SuperSignal West Pico, Thermo Scientific/Pierce, IL, Rockford, USA). For visualization of the signal, the Bio-Rad ChemiDoc Touch Imaging System (Bio-Rad Laboratories, Inc) was used.

Under reducing conditions, the 78 kDa BabA protein band migrated in SDS-PAGE with a molecular mass of ∼75 kDa, whereas it migrated slightly faster under the semi-native conditions as a ∼65 kDa band.

##### SDS-PAGE and immunoblot detection of immune responses from the Leb-mouse vaccination experiment (Figure S4A, B, C, D, E) or semi-native BabA (Figure S4F)

The experimental series was performed similarly to the previous description except that 5% β-mercaptoethanol was used and the bacterial lysate of the challenge onco-strain USU101 was heated by boiling for 5 min. All individual mouse serum samples were analyzed in **Figure S4A, B,** and **C**, whereas seven serum samples were pooled in **Figure S4F**. The immunoblot signal was detected with mouse sera diluted 1:250 and the secondary HRP-conjugated goat anti-mouse antibody diluted 1:1000 (DAKO).

##### Quantification of chemiluminescence signal by Immunoblot scanning, Figure S4D

Images from the ChemiDoc Touch Imaging System (Bio-Rad Laboratories, Inc) were imported into the Bio-Rad image lab software system (Bio-Rad Laboratories, Inc). After the alignment of the image, the bands corresponding to BabA and BabB proteins were searched for in automatic mode. Detected bands were subjected to background substruction and evaluation of the intensity (separately for each sample).

#### 9. 9A *In situ* binding of *H. pylori* to human gastric mucosa histo-tissue sections (Figure 6C)

Bacteria were labelled with fluorescein isothiocyanate (FITC) (Sigma, St. Louis, MO), and *in vitro* bacterial adhesion was tested as described ^44^. Slides were mounted with DAKO fluorescent mounting media (DAKO North America, Inc., CA, USA)) and imaged with a Leica Thunder microscope (Leica Microsystems, Germany). Bacterial attachment was analyzed as described in **ref. Bugaytsova et al., ms1.**

#### 9B. Inhibition of *in situ H. pylori* attachment by mouse sera (Figure 6C)

Inhibition tests were performed with the *in situ* binding methods as described above with the following modifications. FITC-labeled bacteria were first mixed with sera in blocking buffer at a 1:10 dilution on a slowly rocking table for 2 h at room temperature and then processed as described ^44^. Tissue sections of healthy human gastric mucosa were blocked by a mixture of serum from non-infected humans and mice at a final dilution of 1:10. The attachment of bacterial cells was digitalized and quantified as described above except that the quantification was made for the superficial (luminal) epithelium and the bacterial attachment to the gastric pits and glands was excluded.

#### 10. Construction of the phylogenetic tree (Figure S1A)

The phylogenetic tree was calculated based on a core genome alignment of 1,266 genes generated using the Panaroo pan genome pipeline v1.2.10 using 90% protein sequence identity and 75% gene length coverage as the cut-offs ^53,54^. The phylogenetic tree was calculated using PhyML v3.1 ^54^ using the default parameters. To relate CCUG17875 and BCM-300 (GenBank complete assembly GCA_900149805.1) to the worldwide *H. pylori* phylogeography, the following genomic reference sequences were used:

| Genome | GenBank Accession | Hp population |
| --- | --- | --- |
| 26695 | GCA_000008525.1 | hpEurope |
| 51 | GCA_000011725.1 | hpEastAsia |
| 908 | GCA_000148665.1 | hpAfrica1 |
| Aklavik117 | GCA_000315955.1 | hspIndigenousAmerica |
| ausabrJ05 | GCA_001653435.1 | hpSahul |
| F30 | GCA_000270025.1 | hpEastAsia |
| Gambia94/24 | GCA_000185205.1 | hpAfrica1 |
| HPAG1 | GCA_000013245.1 | hpEurope |
| HUP-B14 | GCA_000259235.1 | hpEurope |
| India7 | GCA_000185185.1 | hpAsia2 |
| J99 | GCA_013177275.1 | hpAfrica1 |
| K26A1 | GCA_001653455.1 | hpAfrica2 |
| L7 | GCA_001653375.1 | hpAsia2 |
| NCTC 11637 | GCA_900478295.1 | hpEurope |
| P12 | GCA_000021465.1 | hpEurope |
| PNG84A | GCA_001653475.1 | hpSahul |
| Shi112 | GCA_000277405.1 | hspIndigenousAmerica |
| SouthAfrica7 | GCA_000185245.1 | hpAfrica2 |

#### 11. Quantification and statistical analysis

Data were analyzed using GraphPad Prism 9.0 (Graph Pad software, La Jolla, CA, USA) or R, version 4.0.2 **(R Core Team (2020**). R: A language and environment for statistical computing. R Foundation for Statistical Computing, Vienna, Austria. URL https://www.R-project.org/): Depending on the experimental design and type of variables under investigation, different methods were used. Statistical significance regarding group comparisons was investigated via either unpaired or paired Student’s t-tests or the Wilcoxon signed rank test, while associations between variables were tested using Pearson and/or Spearman correlation. P-values < 0.05 were considered significant (*p < 0.05; **p < 0.01; ***p < 0.001), and p ≥ 0.05 was non-significant (NS). Tests were single-tailed unless otherwise stated.

## Supplemental figures

**Figure S1.**
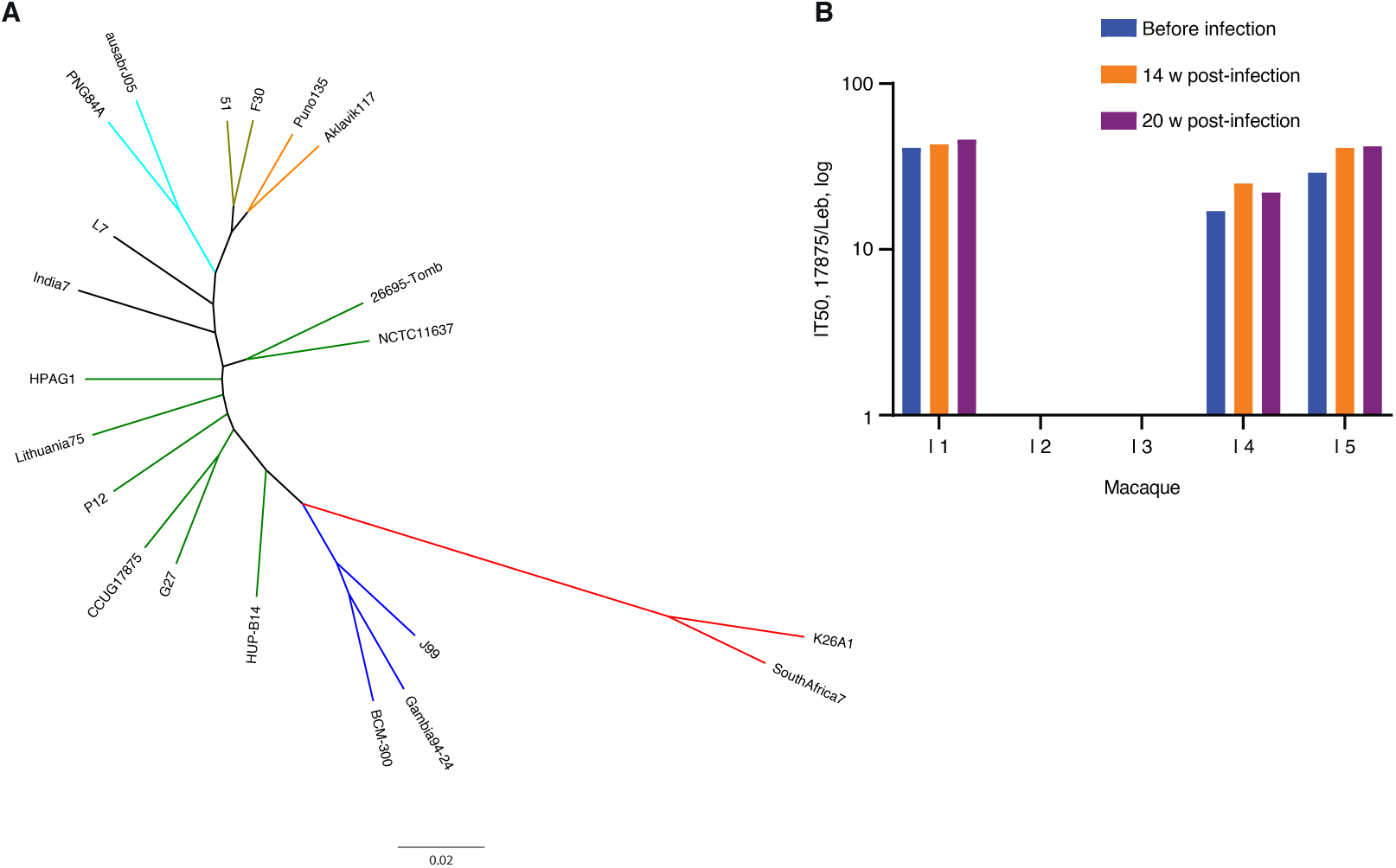
The BCM300 strain phylogeny and induction of IT50 responses by challenge infection in rhesus macaques. **(A)** Maximum likelihood (PhyML) phylogeny relating CCUG 17875 (17875/Leb, GenBank CP090367) and BCM-300 (GenBank assembly GCA_900149805.1) to a reference set of worldwide *H. pylori* genomes. Branch colors denote the *H. pylori* phylogeographic populations hpAfrica2 (Africa, red), hpAfrica1 (Africa blue, including strains BCM-300 and reference strain J99), hpEurope (Europe, green, including reference strains CCUG17875 and 26695-Tomb), hpAsia2 (India, black), hpSahul (Australia and Papua New Guinea, turquoise), hspEAsia (South Kora and Japan, olive), and hspIndigenousAmerica (Peru and Canada, orange). **(B)** Five SPF rhesus macaques were challenge infected with *H. pylori* J166 to test for the induction of IT50 responses. However, three of the animals, no. 1, 4, and 5, demonstrated pre-challenge IT50 titers (in blue) as tested with 17875, suggesting that they carried *H. pylori* before the start of the test.

**Figure S2.**
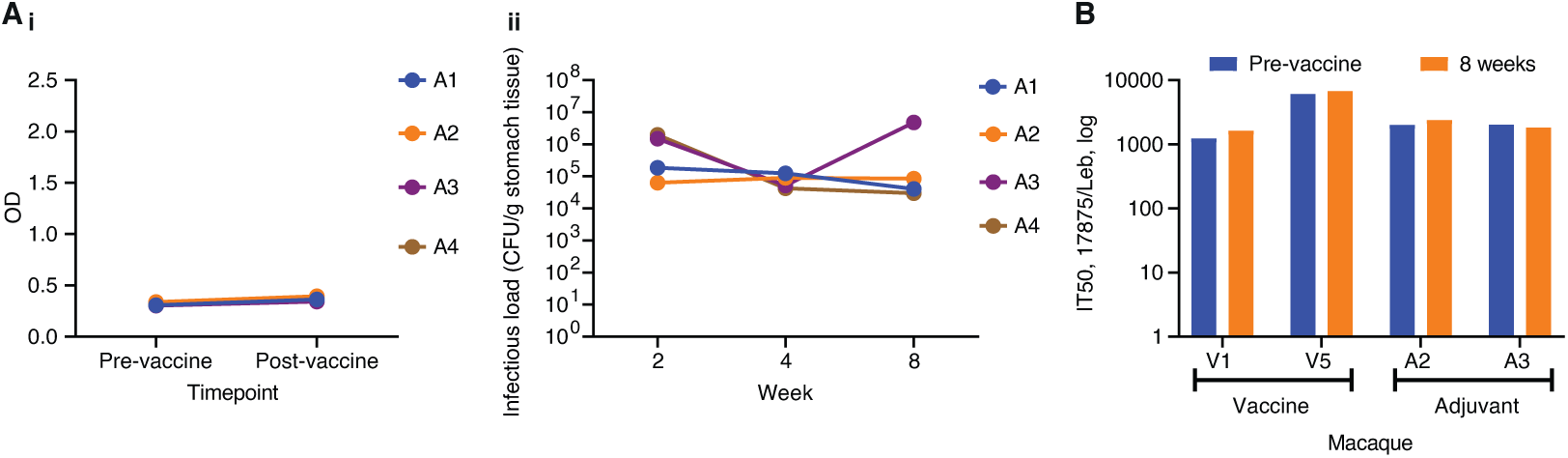
Vaccination and IT50 responses in rhesus macaques. **(A*i*)** ELISA detection of serum BabA antibodies from adjuvant-only control animals, where all four animals were BabA ELISA-negative. **(A*ii*)** The four adjuvant-only control animals demonstrated stable high infection loads during the 8 weeks of infection. **(B)** The vaccinated animals V1 and V5 and the adjuvant-only control animals A2 and A3 demonstrated high pre-challenge IT50 titers when tested with strain 17875.

**Figure S3.**
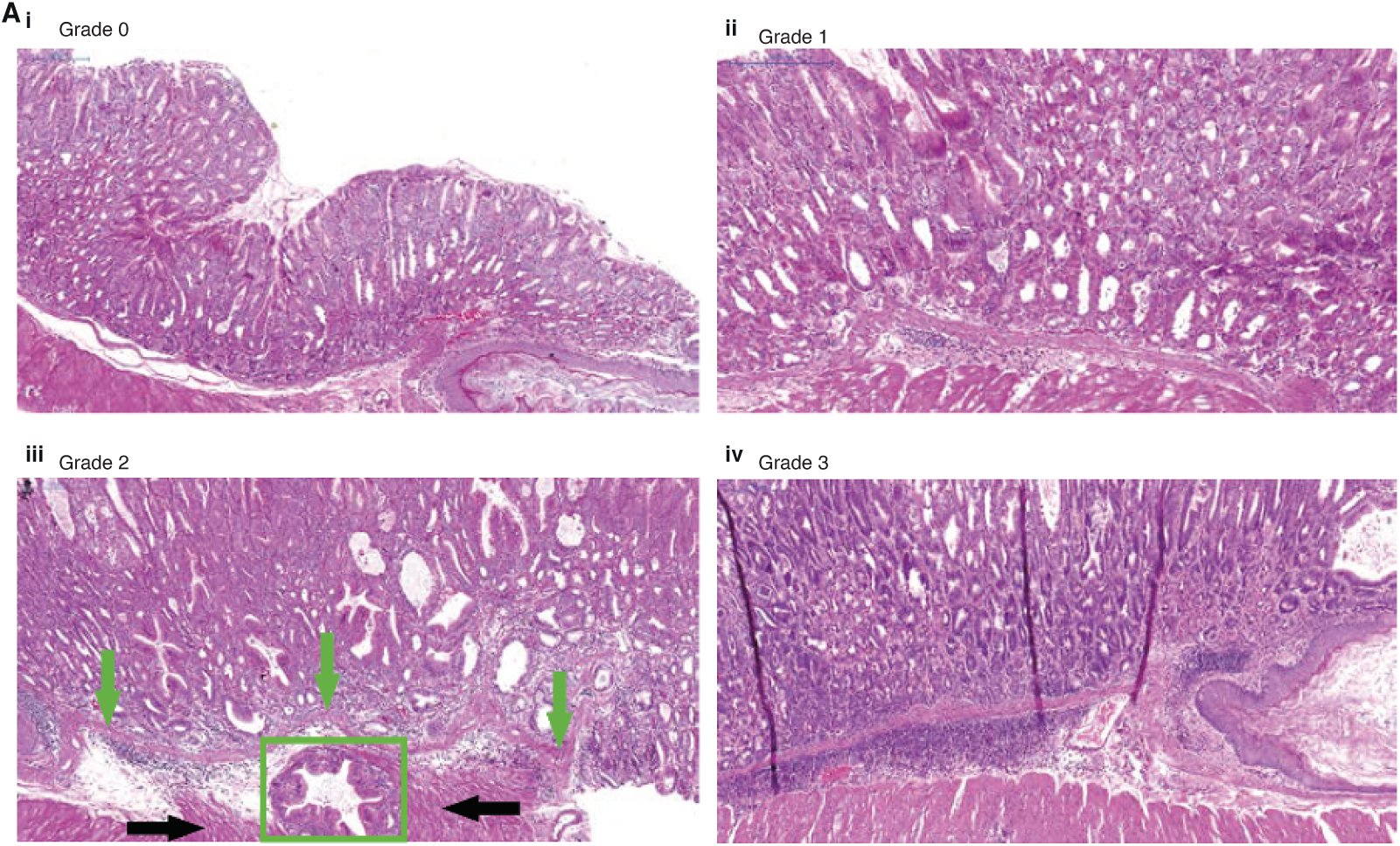
The Leb-mouse gastric cancer animal model. The gastric mucosal inflammatory infiltration scores. **(A*i*)**: Grade 0: The mucous membrane and submucous plate contain rare inflammatory cells, which are diffused throughout the tissue. Inflammatory cells do not form piles (infiltrations) or groups. There are fewer than 20 inflammatory cells (neutrophils, lymphocytes, macrophages, and plasmocytes) visible in the field of view (FOV) (diameter: 1000 µm). **(A*ii*)**: Grade 1: The inflammatory cells are located in the basal part of the mucous membrane and submucous plate where they can form small groups or piles. There are 20–100 inflammatory cells (neutrophils, lymphocytes, macrophages, and plasmocytes) visible in the FOV. **(A*iii*)**: Grade 2: Inflammatory cells are seen in all parts of the mucous membrane and submucous plate. Inflammatory infiltrates (like small lymph nodules) are seen in the basal part of the mucous membrane and submucous plate. The gastric tissue contains 100–300 inflammatory cells per FOV. In addition, the tissue exhibits fulminant gastric adenocarcinoma with growth and penetration through the submucosal layer (lamina propria indicated by green arrows) and with the invasion of cancer tissue (green box) into the muscular layer (indicated by the black arrows). **(A*iv*)**: Grade 3: The inflammatory infiltrates are present in all layers of the gastric tissue and form large inflammatory infiltrates (similar to lymphatic nodes) in the submucous plate (between the muscular and mucus layers). The gastric tissue contains more than 300 inflammatory cells in the FOV.

**Figure S4.**
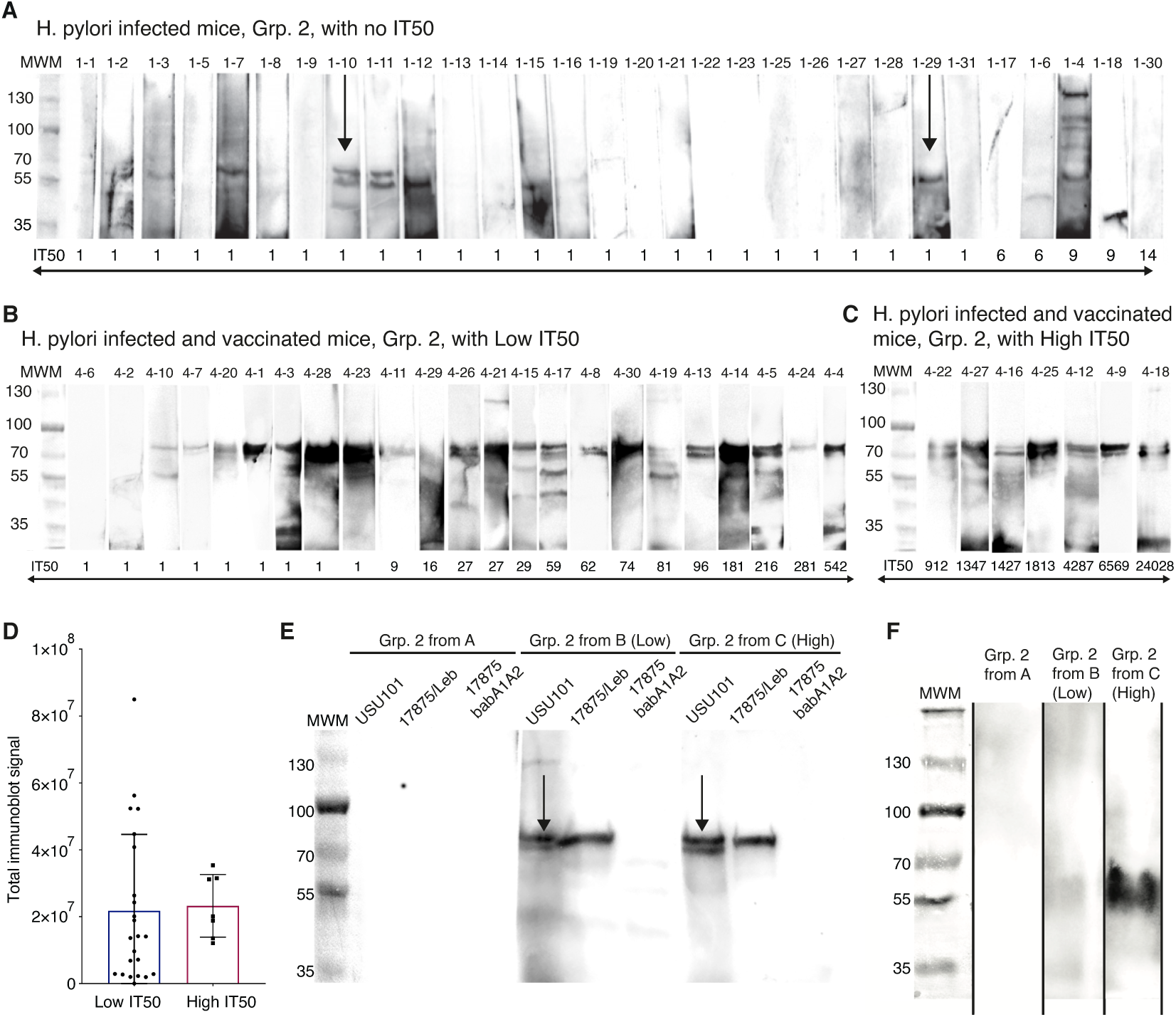
The first vaccination experiment induced protection against inflammation and cancer. The sera samples from the 30 non-vaccinated mice and the 30 vaccinated mice were analyzed by immunoblot detection of whole bacterial protein extracts from *H. pylori* strain USU101, i.e., the strain used for the 12-month infections. The *H. pylori* protein extract was separated on an SDS gel under denaturing (in **A, B, C,** and **E**) or semi-native (**F**) conditions. After immunoblot transfer, the membranes were cut into strips. The immunoblot signals were detected with mouse sera diluted 1:250 and goat-anti-mouse HRP-Ab. The strips in **A, B,** and **C** were arranged according to IT50s. Under reducing conditions, the 78 kDa BabA protein band migrated with a molecular mass of ∼75 kDa; (**A**) with no immuno-detected bands present and in (**B**) and (**C**) where both the BabA and the BabB (migrating slightly faster) bands are present (also in (**E**)). **(A)** The 30 infected but not vaccinated mice all lacked BabA/BabB immunoblot signals, with a median IT50 of 9 ± 5 (similar to the non-infected mice with a median IT50 of 6 ± 5) (**Table 3B** and **3C**). The blot was cut into strips and arranged according to IT50s. **(B)** The 23 infected and vaccinated mice that displayed LOW IT50s with median IT50 = 27 and mean IT50 = 74 (**Table S4A**). The blot was cut into strips and arranged according to IT50s. (C) The seven infected and immunized mice that displayed HIGH IT50s, with median IT50 = 1813 and mean IT50 = 5769 (**Table S4A**). The blot was cut into strips and arranged according to IT50s. **(D)** Scanning of the immunoblot chemiluminescence signal from all immunoblot strips from immunized mice, i.e., from (**B**) with LOW IT50 and from (**C**) with HIGH IT50, showed that the two groups demonstrated no significant difference in intensity in immune response signals against the BabA and BabB bands. Thus, the vaccinated HIGH IT50 mice did not show a stronger signal for linear BabA and BabB epitopes compared to the LOW IT50 mice suggesting that the HIGH IT50 group of mice were not hyper-responding *per se* but rather displayed additional and different immune responses. (**E**) Seven sera samples were pooled from A, from B (Low IT50) and from C (High IT50) and used to probe three immunoblots with whole bacterial protein extracts from strain *H. pylori* USU101 (SabA^−^, BabA^+^, BabB^+^), 17875/Leb (SabA^−^, BabA^+^, BabB^−^), and 17875*bab1babA2* (SabA^+^, BabA^−^, BabB^−^). The *H. pylori* protein extracts were separated on SDS gels under denaturing and reducing conditions. Sera from the infected but non-immunized animals did not display the ∼70–75 kDa BabA/BabB-bands (from **A**), but the bands were visible in sera from immunized animals (from **B** and from **C**) and separated protein extracts from *H. pylori* strains USU101 and 17875/Leb. Two bands were seen on the immunoblot of strain USU101 that expresses both BabA and BabB. In comparison, only the upper ∼75 kDa band was seen for 17875/Leb that expresses BabA but not BabB. In support of this, the ∼70–75 kDa bands were not seen on immunoblot of sera from immunized animals using protein extracts from the *babA*-deletion mutant 17875*bab1babA2* that does not express BabA or BabB. In contrast, the SabA band was not detected in the 17875*bab1babA2* strain, which suggests that SabA is less immunogenic. These results showed that vaccination with BabA and BabB protein raised a humoral immune response against BabA and BabB in almost every vaccinated mouse i.e., the animals in (**B**) and (**C**) with LOW vs. HIGH IT50s. In contrast, the infected but not vaccinated animals (**A**) did not demonstrate natural immune responses towards BabA/BabB. Blots with several dilution series were cut out and aligned for presentation. **(F)** Seven sera samples were pooled from A, from B (Low IT50) and from C (High IT50) and were used to probe three immunoblots with purified BabA protein from strain 17875/Leb. Different from (**E**), in this test the BabA protein was separated by SDS gel electrophoresis under semi-native and non-reducing conditions, similar to Figures 1I and S1H in ***Bugaytsova et al., ms 1*.** The BabA protein was detected only using sera from mice in (**C**), i.e., vaccinated mice with High IT50 responses, whereas mice from groups **A** and **B**, i.e., the infected vs. the infected and vaccinated mice with Low IT50s, provided no immuno-signal with semi-native BabA protein. For this test, the sera samples were 2-fold more dilute (1:500) in order to minimize the (weaker) immune detection of linear epitopes and to preferentially display the (stronger) immune signal of bbAbs that bind to structural and folded BabA epitopes. Thus, the antibodies that were common for both the vaccinated mice with Low (**B**) or High (**C**) IT50s recognize and bind linear BabA epitopes on denaturing immunoblots. In contrast, the bbAbs that were only found in the sera of mice with High IT50 (**C**) can bind to structural BabA epitopes under semi-native conditions. Strips were cut out and aligned because the denatured vs. semi-native samples needed to be separated.

**Figure S5.**
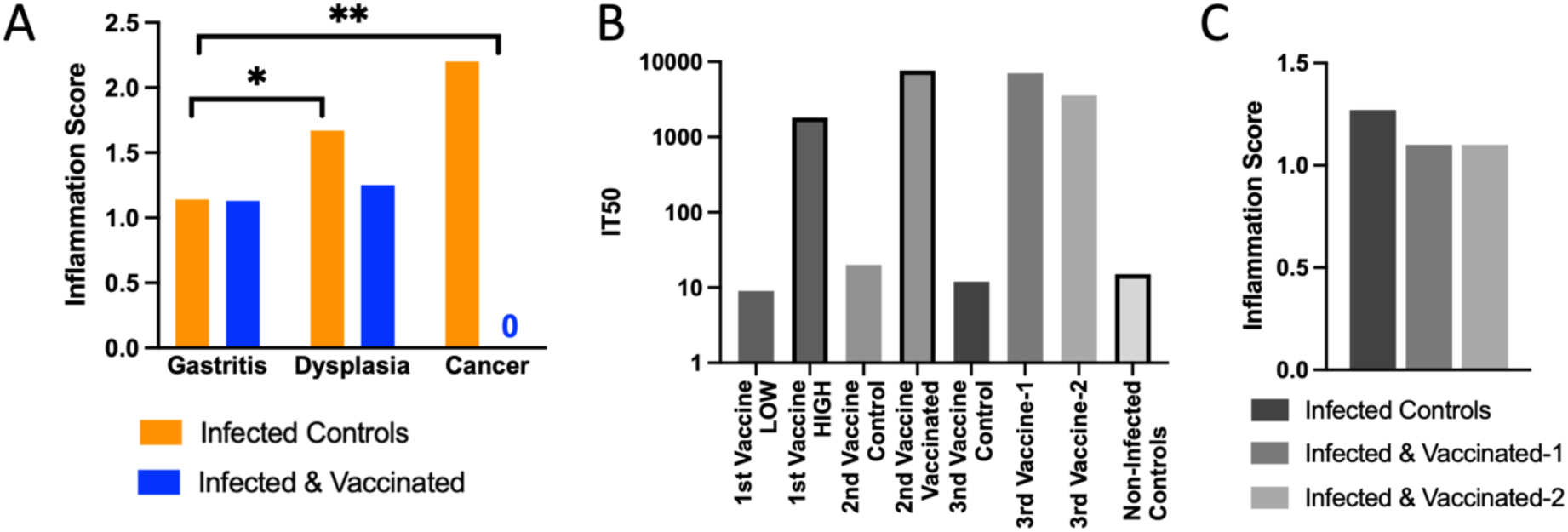
Both the second and third vaccination experiments induced protection against inflammation and cancer. **A**) Similar to the first vaccination experiment, also in the second experimental series the non-vaccinated mice had higher inflammation scores compared to the vaccinated mice. The non-vaccinated mice with dysplasia scored 1.67, Dunn’s test p = 0.039*, and those with cancer scored 2.2, p < 0.001**, compared to the mice with gastritis that scored 1.1. **(B)** The distribution of median IT50s for the first, second, and third vaccine experiments. In the first vaccination experiment, the LOW IT50 group demonstrated a median background IT50 = 16, which was in contrast to the several log-fold higher IT50s, median IT50 = 1813, in the HIGH IT50 group (Figure 4). In the second vaccination experiment (Figures 5 **A-C**), the 16 non-vaccinated control mice demonstrated similar low IT50 = 20, in contrast to the several log-fold higher IT50s, median IT50 = 7318, in the 21 vaccinated mice. Also, in the third vaccination experiment (Figure 5D) the 30 non-vaccinated control mice demonstrated similar low IT50, median IT50 = 15, in contrast to the several log-fold higher IT50s, median IT50 = 7026, in the 27 vaccinated mice in the Vaccine-1 group and the several log-fold higher IT50s, median IT50 = 3557, in the 28 vaccinated mice in the Vaccine-2 group. The 4 non-infected mice demonstrated no or low IT50, with a median IT50 = 15. **(C)** The lower gastric cancer incidence in the third vaccine experiment might be a consequence of the lower mucosal inflammation score of 1.25 in the non-vaccinated animals and 1.1 in the vaccinated animals (Vaccine-1 and Vaccine-2) compared to the higher inflammation score of 1.5 in the non-vaccinated mice in the second vaccine experiment.

**Figure S6.**
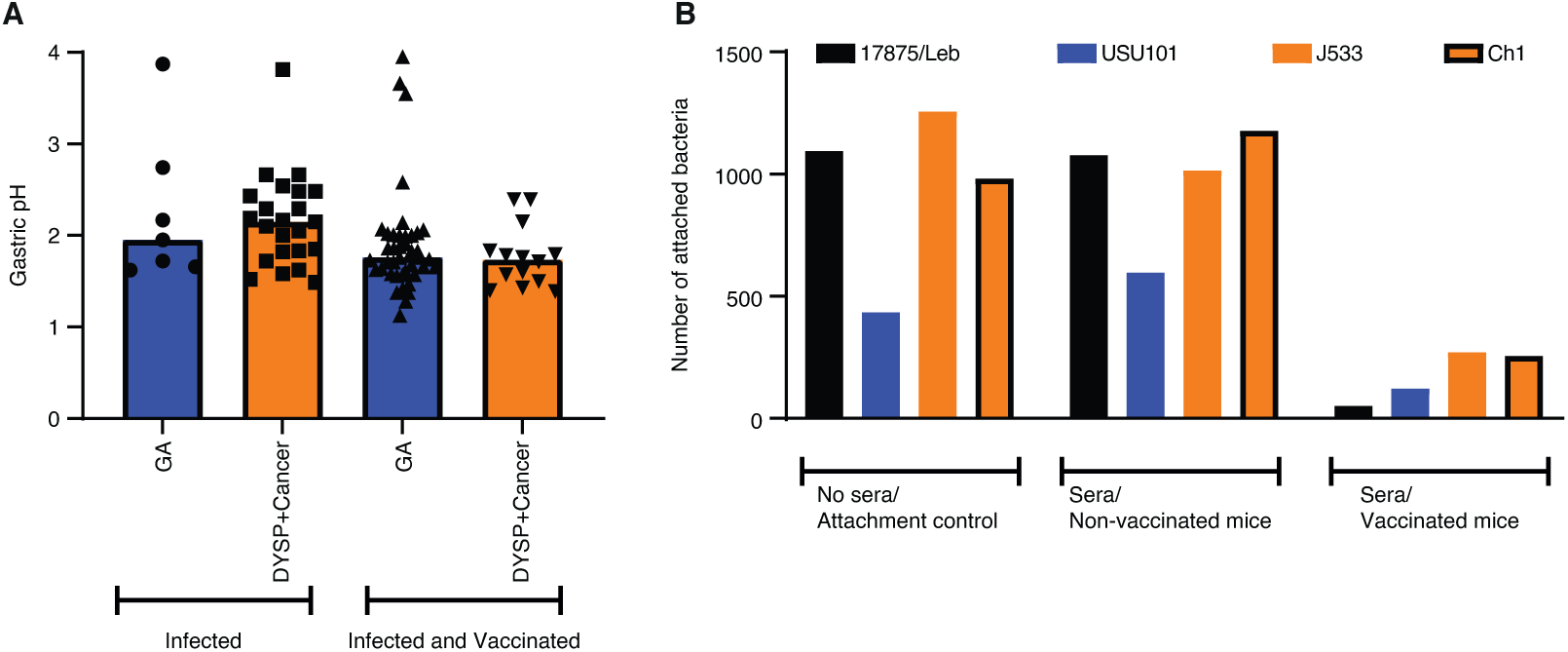
Vaccination preserved gastric acidity and induced bbAb responses that reduced *H. pylori* gastric mucosal attachment. (**A**). In the third vaccination series, the non-vaccinated mice demonstrated elevated gastric pH, whereas the vaccinated groups preserved the gastric acidity over the 12-month period, p = 0.00073***. **(B)** Sera from the second vaccinated group of mice reduced attachment to the human gastric mucosa of *H. pylori* 17875/Leb, USU101, J533 (Japan), and Ch1 (China) compared to the sera of the non-vaccinated controls.

## Supplemental Tables

**Table S1A related to Fig. 1A.** The sera IT50s of healthy volunteers from placebo-controlled phase 1/2 study.

| Serum Code | IT50, *H. pylori* BCM300 |  | IT50, *H. pylori* 17875/Leb |  |
| --- | --- | --- | --- | --- |
|  | Pre-infected | Post-12 weeks | Pre-infected | Post-12 weeks |
| 1001 | 1 | 297 | 1 | 1 |
| 1002 | 1 | 139 | 1 | 1 |
| 1003 | 1 | 267 | 1 | 1 |
| 1004 | 1 | 1823 | 1 | 1 |
| 1006 | 1 | 111 | 1 | 1 |
| 1007 | 1 | 437 | 1 | 30<sup>a</sup> |
| 1008 | 1 | 54 | 1 | 1 |
| 1010 | 1 | 246 | 1 | 1 |
| 1016 | 1 | 181 | 1 | 1 |
| 1018 | 18 | 368 | 1 | 85<sup>a</sup> |
| 1019 | 1 | 1039 | 1 | 1 |
| 1021 | 1 | 285 | 1 | 1 |
| 1022 | 1 | 880 | 1 | 1 |
| 1023 | 1 | 370 | 1 | 1 |
| 1024 | 1 | 1492 | 1 | 51<sup>a</sup> |
| 1027 | 1 | 1 | 1 | 1 |
| 1028 | 1 | 3467 | 1 | 1 |
| 1029 | 1 | 54 | 1 | 16 |
| 1030 | 1 | 2772 | 1 | 27<sup>a</sup> |
| 1031 | 1 | 212 | 1 | 1 |
| 1034 | 1 | 70 | 1 | 1 |
| 1038 | 1 | 128 | 1 | 1 |
| 1039 | 25 | 68 | 1 | 1 |
| 1040 | 1 | 62 | 1 | 1 |
| 1042 | 1 | 58 | 1 | 1 |
| 1047 | 1 | 23 | 1 | 1 |
| 1048 | 1 | 134 | 1 | 50<sup>a</sup> |
| 1049 | 30 | 41 | 1 | 1 |
| 1053 | 1025 | 438 | 1 | 6 |

<sup>a</sup> - Immune response above the critical IT50>30 is indicated by colored boxes.

**Table S1B related to Fig. 1Bi.**
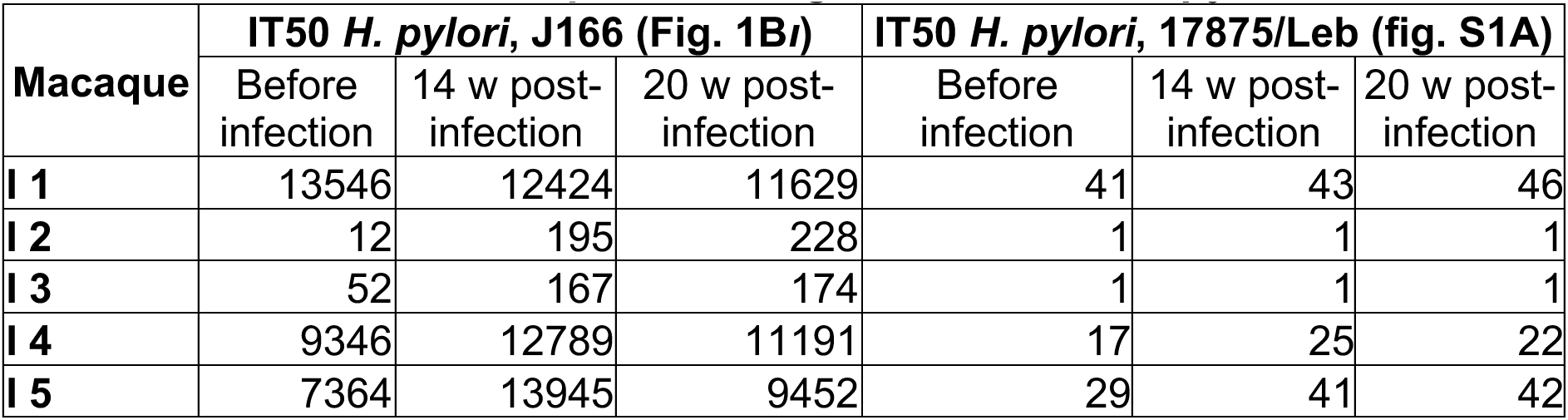
The sera IT50s of SPF macaques challenge infected with *H. pylori* J166 and tested with J166 and 17875/Leb.

**Table S1C related to Fig. 1Bii.** The sera IT50s of SPF macaques challenge infected with *H. pylori* J166 and tested with J166 and I9.

| Macaque | IT50 *H. pylori*, J166 |  | IT50 *H. pylori*, I9 |  |
| --- | --- | --- | --- | --- |
|  | Before infection | 20 w post-infection | Before infection | 20 w post-infection |
| I 2 | 20 | 231 | 16 | 45 |
| I 3 | 65 | 223 | 32 | 50 |

**Table S2A related to Fig. 2Ai and fig. S2 Ai.** The sera ELISA of vaccinated and control macaques.

| Time-point | ELISA |  |  |  |  |  |  |  |  |
| --- | --- | --- | --- | --- | --- | --- | --- | --- | --- |
|  | V1<sup>a</sup> | V2 | V3 | V4 | V5 | A1<sup>b</sup> | A2 | A3 | A4 |
| Pre-vaccine | 0.336 | 0.351 | 0.361 | 0.366 | 0.282 | 0.307 | 0.339 | 0.303 | 0.318 |
| Postvaccine | 0.501 | 1.950 | 2.371 | 0.950 | 1.382 | 0.363 | 0.395 | 0.343 | 0.375 |

V<sup>a</sup> – vaccinated macaque
A<sup>b</sup> – control macaque with adjuvant only

**Table S2B related to Fig. 2Aii and Fig. S2Aii.** *H. pylori* infectious load in vaccinated and control macaques.

| Time after vaccination | Infectious load (CFU/g of stomach tissue) |  |  |  |  |  |  |  |  |
| --- | --- | --- | --- | --- | --- | --- | --- | --- | --- |
|  | V1<sup>a</sup> | V2 | V3 | V4 | V5 | A1<sup>b</sup> | A2 | A3 | A4 |
| Week 2 | 95.500 | 140 | 665 | 2.750.000 | 132.000 | 187.000 | 63.900 | 1.500.000 | 1.970.000 |
| Week 4 | 36.100 | 1 | 1 | 2.880.000 | 383.000 | 125.000 | 90.900 | 52.400 | 42.900 |
| Week 8 | 154.000 | 205 | 6.290 | 40.600.000 | 2.410.000 | 40.500 | 85.300 | 4.860.000 | 29.900 |

V<sup>a</sup> – vaccinated macaque
A<sup>b</sup> – control macaque with adjuvant only

**Table S2C related to Fig. 2B.** The sera IT50s of vaccinated and control macaques.

| Time-point | IT50, *H. pylori* 17875/Leb and I9 |  |  |  |  |  |  |  |  |  |
| --- | --- | --- | --- | --- | --- | --- | --- | --- | --- | --- |
|  | V2<sup>a</sup> |  | V3 |  | V4 |  | A1<sup>b</sup> |  | A4 |  |
|  | 17875/Leb | I9 | 17875/Leb | I9 | 17875/Leb | I9 | 17875/Leb | I9 | 17875/Leb | I9 |
| Pre-serum | 4 | 18 | 1 | 38 | 1 | 7 | 1 | 30 | 1 | 17 |
| Week 2 | 8 | 63 | 30 | 102 | 17 | 58 | 1 | 30 | 1 | 25 |
| Week 4 | 25 | 79 | 28 | 131 | 18 | 52 | 1 | - | 1 | - |
| Week 8 | 46 | 89 | 17 | 171 | 12 | 56 | 1 | - | 1 | - |

V<sup>a</sup> – vaccinated macaque
A<sup>b</sup> – control macaque with adjuvant only

**Table S3A related to Fig 3B.** Sera IT50 for *H. pylori* 17875/Leb and Histological Scoring of *H. pylori* infected and vaccinated Leb-mice. Group I, Infected Mice (34 mice).

| Mouse | CFU/g | Leb binding, % | Chronic Gastritis | Dysplasia | Cancer | Inflammation Score | Final Diagnosis | IT50 |
| --- | --- | --- | --- | --- | --- | --- | --- | --- |
| #1-1 | 13333 | 0 | + | + | + | 2 | Cancer | 1 |
| #1-2 | 0 | - | + | - | - | 1 | GA | 1 |
| #1-4 | 1600 | 10 | + | + | - | 1 | DYSP | 1 |
| #1-5 | 1278 | 0 | + | - | - | 2 | GA | 1 |
| #1-8 | 0 | - | + | + | + | 3 | Cancer | 1 |
| #1-10 | 0 | - | + | + | + | 3 | Cancer | 1 |
| #1-12 | 0 | - | + | - | - | 2 | GA | 1 |
| #1-13 | 0 | - | + | - | - | 1 | GA | 1 |
| #1-14 | 6321 | 0 | + | + | + | 3 | Cancer | 1 |
| #1-15 | 438 | 0 | + | + | + | 2 | Cancer | 1 |
| #1-16 | 0 | - | + | - | - | 1 | GA | 1 |
| #1-18 | 469 | 0 | + | + | + | 3 | Cancer | 1 |
| #1-19 | 0 | - | + | + | + | 2 | Cancer | 1 |
| #1-21 | 125 | 0 | + | + | + | 3 | Cancer | 1 |
| #1-24 | 0 | - | + | + | - | 1 | DYSP | 1 |
| #1-25 | 3950 | 0 | + | + | + | 3 | Cancer | 1 |
| #1-27 | 0 | - | + | - | - | 1 | GA | 1 |
| #1-28 | 39 | 0 | + | + | + | 3 | Cancer | 1 |
| #1-29 | 563 | 0 | + | + | + | 3 | Cancer | 1 |
| #1-30 | 0 | - | + | - | - | 2 | GA | 1 |
| #1-31 | 145 | 0 | + | + | + | 3 | Cancer | 1 |
| #1-32 | 886 | 0 | + | + | + | 3 | Cancer | 1 |
| #1-33 | 283 | 0 | + | + | + | 3 | Cancer | 1 |
| #1-34 | 0 | - | + | + | + | 3 | Cancer | 1 |
| #1-35 | 205 | 0 | + | + | - | 2 | DYSP | 2 |
| #1-17 | 2200 | 0 | + | - | - | 2 | GA | 3 |
| #1-6 | 0 | - | + | + | + | 3 | Cancer | 3 |
| #1-7 | 0 | -- | + | - | - | 1 | GA | 4 |
| #1-20 | 0 | - | + | + | + | 3 | Cancer | 6 |
| #1-26 | 0 | - | + | + | + | 3 | Cancer | 7 |
| #1-3 | 0 | - | + | + | - | 2 | DYSP | 7 |
| #1-23 | 0 | - | + | + | - | 2 | DYSP | 7 |
| #1-22 | 0 | - | + | + | - | 2 | DYSP | 7 |
| #1-9 | 4556 | 0 | + | + | + | 3 | Cancer | 10 |
|  | 16 CFU positive/34 mice |  |  |  |  | 77/34 animals=2.26 |  |  |
|  | Mean, 2274 |  |  |  |  |  |  |  |
|  | Median, 725 |  |  |  |  |  |  |  |

**Table S3B related to Fig 3B.** Group II, Infected Mice (30 mice).

| Mouse | CFU/g | Leb binding, % | Chronic Gastritis | Dysplasia | Cancer | Inflammation Score | Final Diagnosis | IT50 |
| --- | --- | --- | --- | --- | --- | --- | --- | --- |
| #1-1 | 4222 | 0 | + | - | - | 1 | GA | 1 |
| #1-2 | 0 | - | + | - | - | 0 | GA | 1 |
| #1-3 | 0 | - | + | - | - | 0 | GA | 1 |
| #1-5 | 0 | - | + | + | + | 3 | Cancer | 1 |
| #1-7 | 1000 | 0 | + | + | + | 2 | Cancer | 1 |
| #1-8 | 0 | - | + | + | - | 0 | DYSP | 1 |
| #1-9 | 200 | 3 | + | + | - | 1 | DYSP | 1 |
| #1-10 | 263 | 0 | + | + | + | 3 | Cancer | 1 |
| #1-11 | 8499 | 0 | + | + | + | 3 | Cancer | 1 |
| #1-12 | 10500 | 0 | + | + | + | 3 | Cancer | 1 |
| #1-13 | 0 | - | + | - | - | 1 | GA | 1 |
| #1-14 | 0 | - | + | - | - | 0 | GA | 1 |
| #1-15 | 0 | - | + | - | - | 1 | GA | 1 |
| #1-16 | 0 | - | + | + | + | 2 | Cancer | 1 |
| #1-19 | 0 | - | + | + | + | 3 | Cancer | 1 |
| #1-20 | 0 | - | + | + | - | 2 | DYSP | 1 |
| #1-21 | 80 | 0 | + | + | - | 1 | DYSP | 1 |
| #1-22 | 0 | - | + | - | - | 1 | GA | 1 |
| #1-23 | 0 | - | + | - | - | 1 | GA | 1 |
| #1-25 | 0 | - | + | - | - | 0 | GA | 1 |
| #1-26 | 0 | - | + | + | - | 0 | DYSP | 1 |
| #1-27 | 0 | - | + | + | - | 1 | DYSP | 1 |
| #1-28 | 0 | - | + | + | - | 1 | DYSP | 1 |
| #1-29 | 947 | 0 | + | + | + | 2 | Cancer | 1 |
| #1-31 | 0 | - | + | + | + | 1 | Cancer | 1 |
| #1-6 | 0 | - | + | + | + | 1 | Cancer | 6 |
| #1-17 | 0 | - | + | + | - | 2 | DYSP | 6 |
| #1-4 | 57629 | 0 | + | + | + | 3 | Cancer | 9 |
| #1-18 | 0 | - | + | - | - | 1 | GA | 9 |
| #1-30 | 0 | - | + | + | + | 2 | Cancer | 14 |
|  | 9 CFU positive/30 mice |  |  |  |  | 42/30 animals=1.4 |  |  |
|  | Mean, 9260 |  |  |  |  |  |  |  |
|  | Median, 1000 |  |  |  |  |  |  |  |

**Table S3C related to Fig 3B.**
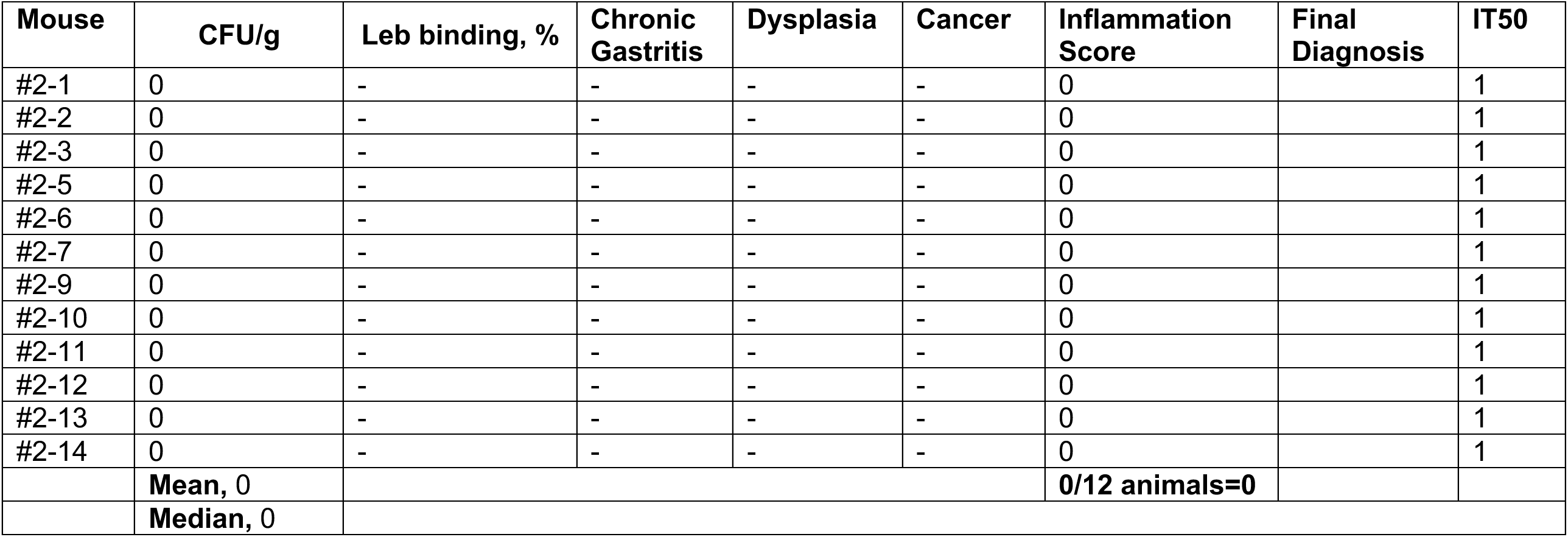
Group III, Non-infected Mice (12 mice).

**Table S3D related to Fig. 3C.**
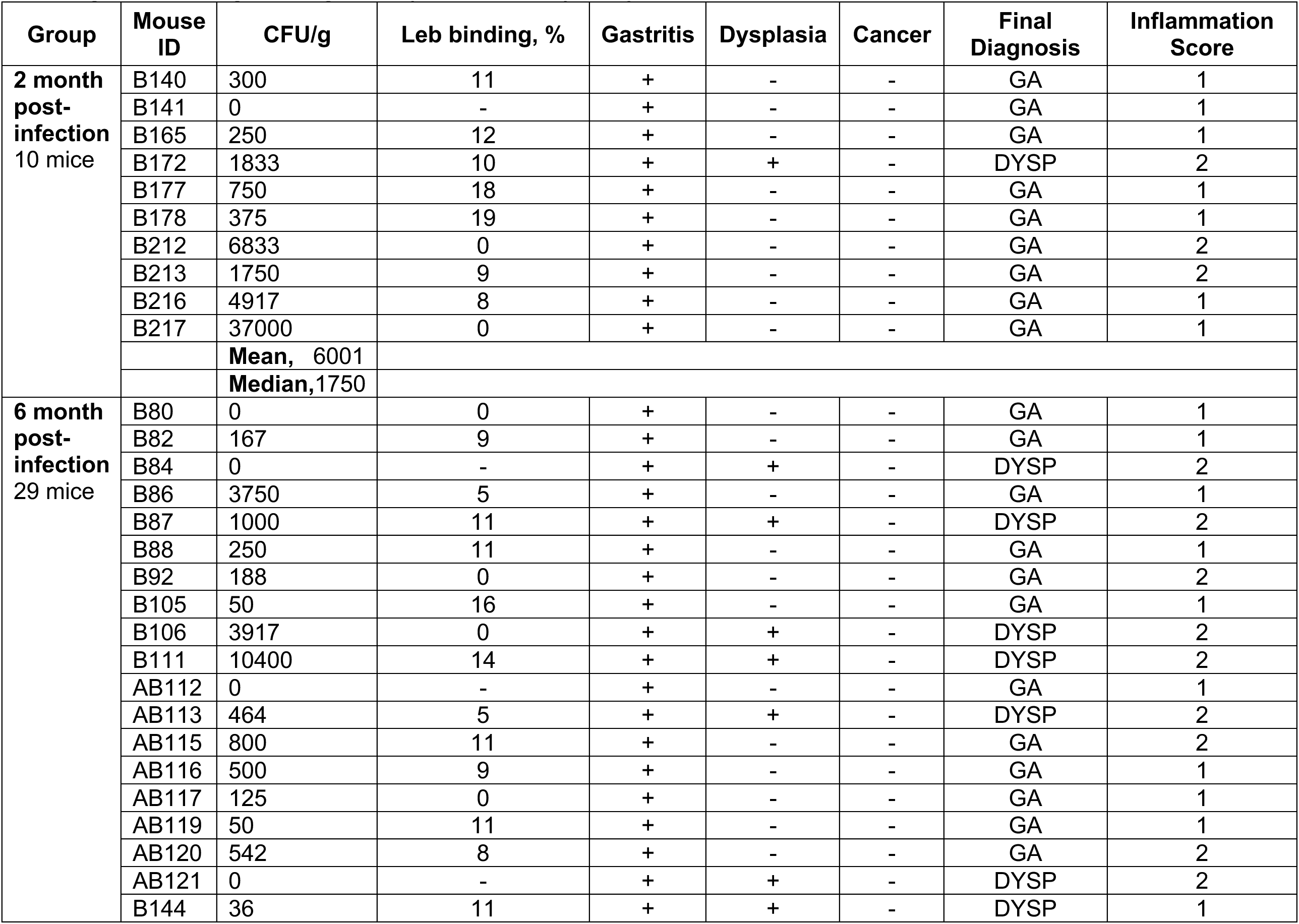
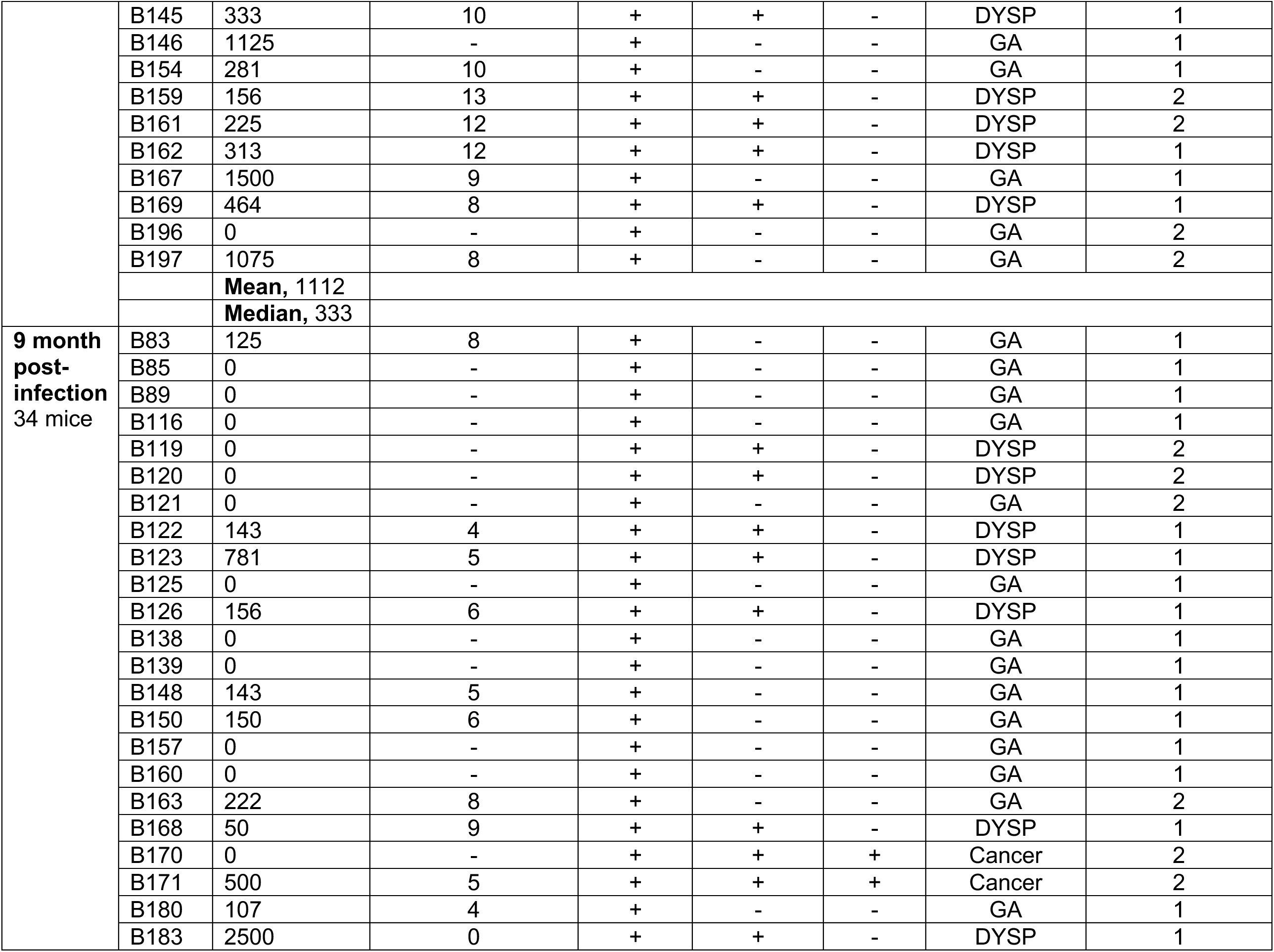
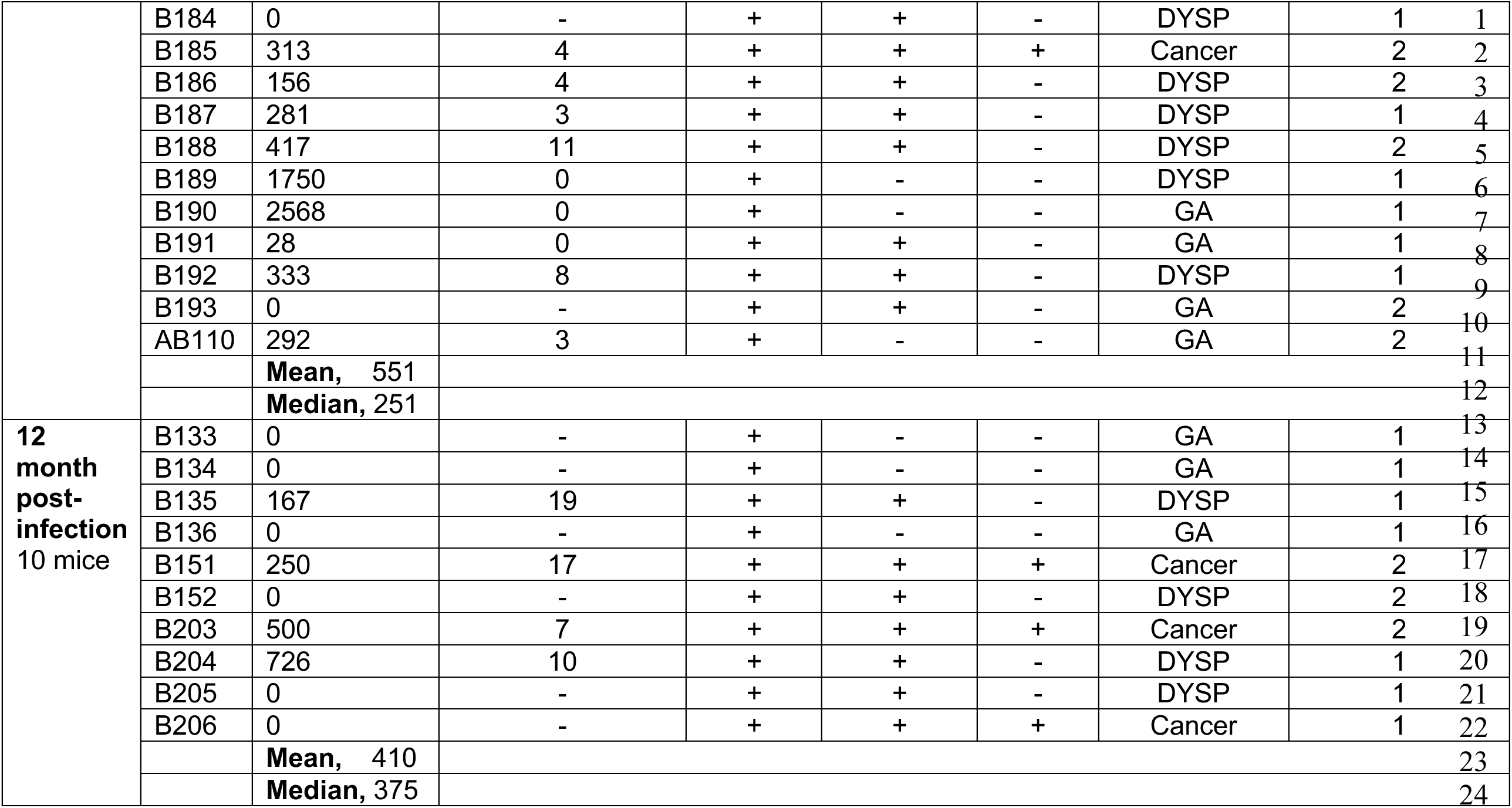
Histological Scoring of long-term (2-12 months) *H. pylori* infected Leb-mice.

**Table S3E related to Fig. 3D.**
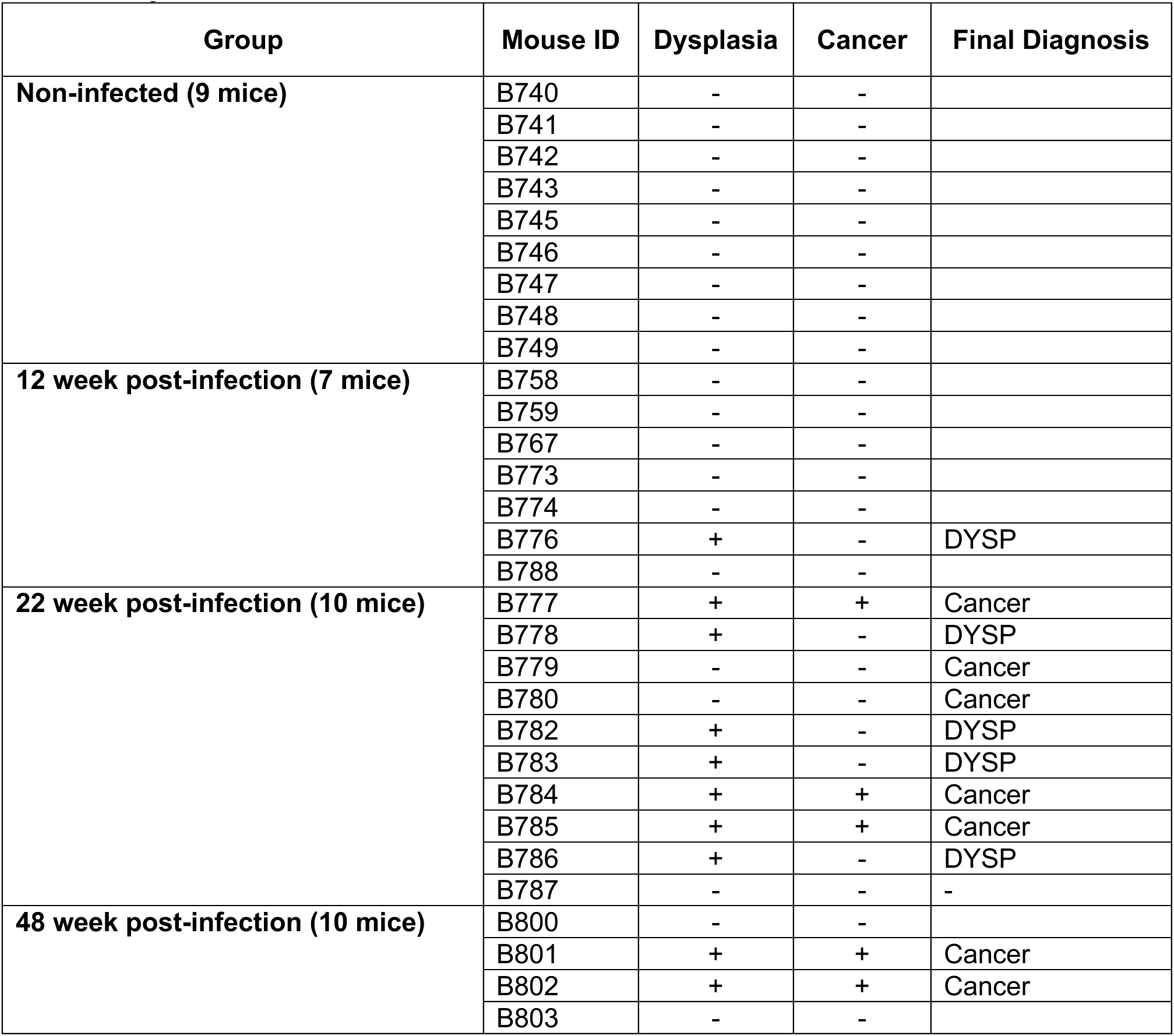
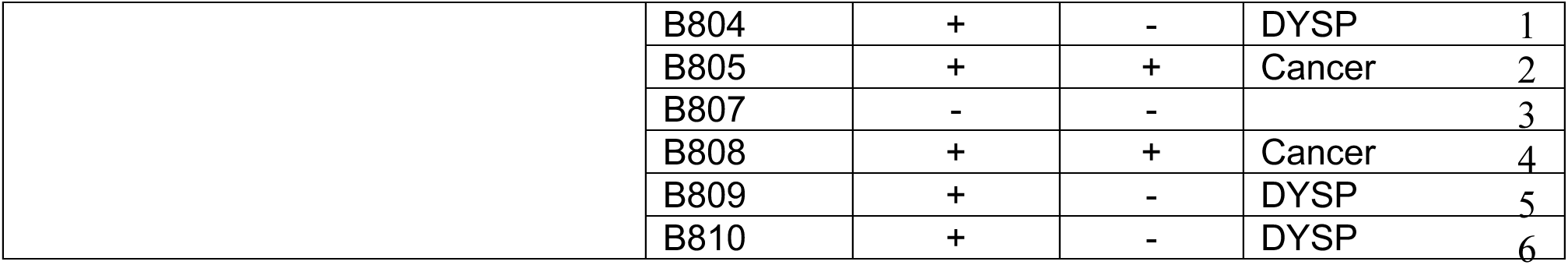
Sera IT50 for *H. pylori* 17875/Leb and Histological Scoring of *H. pylori* infected Leb-mice after eradiction of infection by antibiotics.

| Group | Mouse ID | Dysplasia | Cancer | Final Diagnosis |
| --- | --- | --- | --- | --- |
| Non-infected (9 mice) | B740 | - | - |  |
|  | B741 | - | - |  |
|  | B742 | - | - |  |
|  | B743 | - | - |  |
|  | B745 | - | - |  |
|  | B746 | - | - |  |
|  | B747 | - | - |  |
|  | B748 | - | - |  |
|  | B749 | - | - |  |
| 12 week post-infection (7 mice) | B758 | - | - |  |
|  | B759 | - | - |  |
|  | B767 | - | - |  |
|  | B773 | - | - |  |
|  | B774 | - | - |  |
|  | B776 | + | - | DYSP |
|  | B788 | - | - |  |
| 22 week post-infection (10 mice) | B777 | + | + | Cancer |
|  | B778 | + | - | DYSP |
|  | B779 | - | - | Cancer |
|  | B780 | - | - | Cancer |
|  | B782 | + | - | DYSP |
|  | B783 | + | - | DYSP |
|  | B784 | + | + | Cancer |
|  | B785 | + | + | Cancer |
|  | B786 | + | - | DYSP |
|  | B787 | - | - | - |
| 48 week post-infection (10 mice) | B800 | - | - |  |
|  | B801 | + | + | Cancer |
|  | B802 | + | + | Cancer |
|  | B803 | - | - |  |

|  |  |  |  |  |  |
| --- | --- | --- | --- | --- | --- |
|  | B804 | + | - | DYSP | 1 |
|  | B805 | + | + | Cancer | 2 |
|  | B807 | - | - |  | 3 |
|  | B808 | + | + | Cancer | 4 |
|  | B809 | + | - | DYSP | 5 |
|  | B810 | + | - | DYSP | 6 |

**Table S3F related to Figure 3E and 3F.** Prevalence of CFU-positive *H. pylori* Leb-mice and their Leb-binding activity at the 12 months end-point.

| Group | Treatment | CFU-positive mice, % | Leb-binders among outputs, % |
| --- | --- | --- | --- |
| Group I, Fig. 3B | *H. pylori* infected Leb mice | 47 | 6 |
| Group II, Fig. 3B | *H. pylori* infected Leb mice | 30 | 11 |
| Group Infected/Controls, Fig. 5 ABC | *H. pylori* infected Leb mice | 38 | 16 |
| Group Infected/Controls, Fig.5 D | *H. pylori* infected Leb mice | 20 | 66 |
| Group 12m, Fig. 3C | *H. pylori* infected Leb mice | 40 | 40 |
|  | **Median** | **38** | **16** |
|  | **Mean** | **35** | **28** |

**Table S4A. related to Fig. 4B.** Group II, Infected and Vaccinated LOW and HIGH IT50 Mice (30 mice).

| Mouse | CFU/g | Leb binding, % | Chronic Gastritis | Dysplasia | Cancer | Inflammation Score | Final Diagnosis | IT50 |
| --- | --- | --- | --- | --- | --- | --- | --- | --- |
| **Low IT50 (23 mice)** |  |  |  |  |  |  |  |  |
| #4-1 | 0 | - | + | + | - | 0 | DYSP | 1 |
| #4-2 | 1571 | 0 | + | + | + | 3 | Cancer | 1 |
| #4-3 | 0 | - | + | + | + | 3 | Cancer | 1 |
| #4-6 | 0 | - | + | + | - | 1 | DYSP | 1 |
| #4-7 | 0 | - | + | + | - | 1 | DYSP | 1 |
| #4-10 | 0 | - | + | - | - | 0 | GA | 1 |
| #4-20 | 441 | 5 | + | + | - | 2 | DYSP | 1 |
| #4-23 | 0 | - | + | - | - | 1 | GA | 1 |
| #4-28 | 200 | 0 | + | + | - | 0 | DYSP | 1 |
| #4-11 | 0 | - | + | - | - | 1 | GA | 9 |
| #4-29 | 800 | 0 | + | + | + | 2 | Cancer | 16 |
| #4-21 | 1312 | 0 | + | + | + | 3 | Cancer | 27 |
| #4-26 | 0 | - | + | + | + | 2 | Cancer | 27 |
| #4-15 | 100 | 10 | + | + | + | 2 | Cancer | 29 |
| #4-17 | 0 | - | + | + | + | 2 | Cancer | 59 |
| #4-8 | 0 | - | + | + | + | 1 | Cancer | 62 |
| #4-30 | 400 | 9 | + | + | + | 1 | Cancer | 74 |
| #4-19 | 3583 | 0 | + | + | + | 2 | Cancer | 81 |
| #4-13 | 0 | - | + | + | + | 3 | Cancer | 96 |
| #4-14 | 0 | - | + | + | + | 3 | Cancer | 181 |
| #4-24 | 0 | - | + | + | + | 1 | Cancer | 281 |
| #4-5 | 4317 | 0 | + | + | + | 3 | Cancer | 216 |
| #4-4 | 0 | - | + | + | + | 2 | Cancer | 542 |
|  |  |  |  |  |  | 39/23 animals=1.7 |  | Median, 27 |
| **High IT50 (7 mice)** |  |  |  |  |  |  |  |  |
| #4-22 | 0 | - | + | - | - | 0 | GA | 912 |
| #4-27 | 1500 | 0 | + | + | + | 2 | Cancer | 1347 |
| #4-16 | 0 | - | + | - | - | 1 | GA | 1427 |
| #4-25 | 0 | - | + | - | - | 0 | GA | 1813 |
| #4-12 | 1466 | 0 | + | + | - | 2 | DYSP | 4287 |
| #4-9 | 0 | - | + | + | + | 2 | Cancer | 6569 |
| #4-18 | 0 | - | + | - | - | 0 | GA | 24028 |
|  |  |  |  |  |  | 46/30 animals=1.5 |  | Median, 1813 |

**Table S4B related to Fig. 4C.** The 1st Vaccine Experiment. Summary of incidense of disease. The panel shows the number of animals in the group of “Infected and vaccinated” with a 3.0 *vs*. 0.5 ratio of Cancer/Gastritis for the Low (23 animals) *vs.* High (7 animals) IT50 groups, respectively. The difference in ratios demonstrated a 6-fold reduced risk for gastric cancer in the High IT50 group (p < 0.038).

| Titer | GAstritis | DYSPlasia | Cancer | <u>Cancer</u><br>GAstritis |
| --- | --- | --- | --- | --- |
| Low IT50 (L) | 3 | 5 | 15 | 3 |
| High IT50 (H) | 4 | 1 | 2 | 0.5 |
| Ratio Low IT50 / High IT50 |  |  |  | 6 |
| Fisher's Test |  |  |  | 0.038 |

**Table S4C related to Fig. 3 and Fig. 4.**
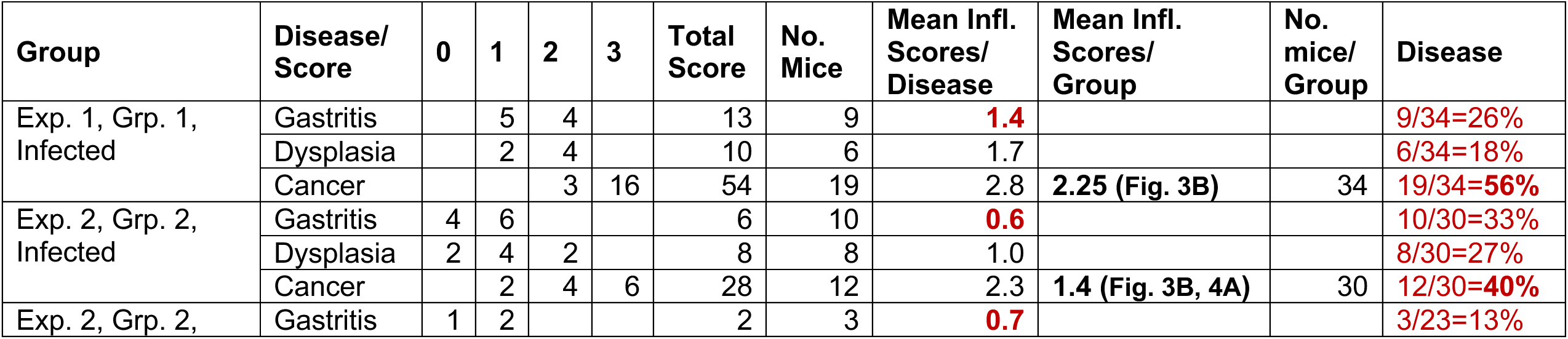
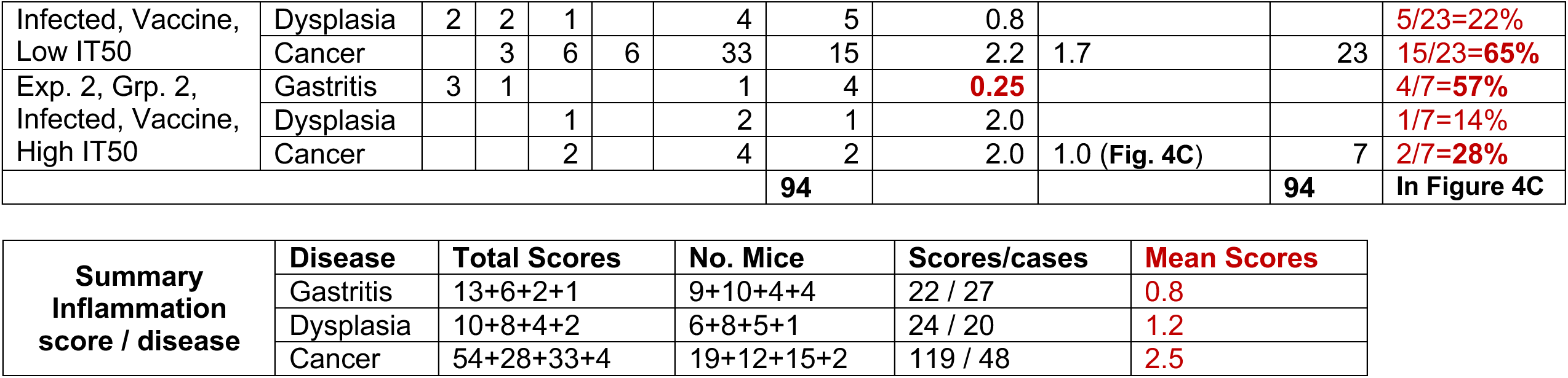
The Cancer model Summary of disease and inflammation scores.

**Table S4D related to Fig. 5.** The 2^nd^ Vaccine Experiment related to Fig. 5ABC.

| Mouse | CFU/g | Leb binding, % | Gastric pH | Chronic Gastritis | Dysplasia | Cancer | Inflammation Score | Final Diagnosis | IT50, 17875/Leb |
| --- | --- | --- | --- | --- | --- | --- | --- | --- | --- |
| **Infected mice (16 mice)** |  |  |  |  |  |  |  |  |  |
| B891 |  |  | 1.5 | + | - | - | 1 | GA | 20 |
| B892 |  |  | 2.37 | + | + | - | 2 | DYSP | 16 |
| B893 | 7700 | 3 | 1.77 | + | + | + | 3 | Cancer | 22 |
| B894 | 56 | X | 2.8 | + | + | + | 2 | Cancer | 10 |
| B895 |  |  | 1.54 | + | + | - | 2 | DYSP | 6 |
| B896 |  |  | 4.44 | + | - | - | 1 | GA | 24 |
| B897 | 750 | 0 | 3.19 | + | + | - | 1 | DYSP | 24 |
| B898 |  |  | 1.92 | - | - | - | 0 | Healthy | 20 |
| B912 |  |  | 3.04 | + | + | + | 1 | Cancer | 17 |
| B922 |  |  | 1.82 | + | + | + | 2 | Cancer | 10 |
| B935 | 813 | X | 1.27 | + | - | - | 2 | GA | 22 |
| B936 | 929 | 0 | 3.26 | + | + | + | 3 | Cancer | 24 |
| B937 |  |  | 1.52 | + | - | - | 1 | GA | 13 |
| B938 |  |  | 1.35 | + | - | - | 1 | GA | 30 |
| B940 |  |  | 1.33 | + | - | - | 1 | GA | 8 |
| B942 | 167 | X | 1.34 | + | - | - | 1 | GA | 21 |
|  |  |  |  |  |  |  | Mean, 1.5 |  | Median, 20 |
| **Infected and Vaccinated mice (21 mice)** |  |  |  |  |  |  |  |  |  |
| B899 |  |  | 2.4 | + | - | - | 1 | GA | 7667 |
| B900 |  |  | 4.96 | + | - | - | 1 | GA | 8539 |
| B901 |  |  | 3.53 | - | - | - | 0 | Healthy | 8698 |
| B903 |  |  | 1.82 | + | - | - | 1 | DYSP | 33622 |
| B904 | 150 | 16 | 3.83 | + | - | - | 1 | GA | 3959 |
| B905 |  |  | 1.77 | + | - | - | 1 | GA | 4146 |
| B906 | 2107 | 3 | 1.66 | + | - | - | 1 | GA | 3969 |
| B907 | 143 | 0 | 1.47 | + | - | - | 2 | GA | 6163 |
| B908 | 458 | 7 | 1.79 | + | - | - | 1 | GA | 2270 |
| B909 |  |  | 1.7 | + | - | - | 1 | GA | 7318 |
| B910 |  |  | 2.18 | + | - | - | 1 | GA | 764 |
| B917 | 286 | 1 | 1.86 | + | - | - | 0 | Healthy | 12035 |
| B918 |  |  | 1.64 | + | - | - | 1 | GA | 11284 |
| B924 |  |  | 2.48 | + | + | - | 1 | DYSP | 5798 |
| B927 |  |  | 1.33 | + | - | - | 1 | GA | 5663 |
| B929 |  |  | 1.49 | + | + | - | 2 | DYSP | 10934 |
| B930 |  |  | 1.68 | + | - | - | 2 | GA | 13338 |
| B931 | 333 | X | 1.87 | + | - | - | 1 | GA | 33113 |
| B932 |  |  | 1.43 | + | + | - | 1 | DYSP | 40035 |
| B933 | 50 | X | 1.45 | + | - | - | 1 | GA | 12999 |
| B934 | 83 | X | 1.34 | + | - | - | 1 | GA | 5893 |
|  |  |  |  |  |  |  | Mean, **1.05** |  | Median, 7667 |

1 X – slow growing outputs

**Table S4E.** The 3^rd^ Vaccine Experiment related to Fig. 5D. **Infected (30 mice) and Vaccinated w High antigen dose (27 mice) and, Infected and Vaccinated with Low antigen dose (28 mice)**

| Mouse | CFU/g | Leb binding, % | Gastric pH | Chronic Gastritis | Dysplasia | Cancer | Inflammation Score | Final Diagnosis |
| --- | --- | --- | --- | --- | --- | --- | --- | --- |
| **Infected control mice (30 mice)** |  |  |  |  |  |  |  |  |
| B988 |  |  | 2.19 | + | + | + | 2 | Cancer |
| B989 | 267 | 15 | 2.17 | + | + | - | 2 | DYSP |
| B990 |  |  | 2.54 | + | + | + | 3 | Cancer |
| B991 |  |  | 3.87 | + | - | - | 1 | GA |
| B992 | 154 | 0 | 1.62 | + | + | - | 1 | DYSP |
| B993 |  |  | 1.62 | + | - | - | 1 | GA |
| B994 |  |  | 1.82 | + | + | - | 1 | DYSP |
| B995 |  |  | 1.72 | + | - | - | 1 | GA |
| B1013 |  |  | 2.17 | + | - | - | 1 | GA |
| B1014 |  |  | 1.95 | + | - | - | 1 | GA |
| B1015 |  |  | 1.49 | + | + | - | 2 | DYSP |
| B1016 |  |  | 2.29 | + | - | - | 1 | GA |
| B1017 |  |  | 2.05 | + | + | + | 1 | Cancer |
| B1018 |  |  | 2.66 | + | - | - | 2 | GA |
| B1019 |  |  | 2.48 | + | + | + | 1 | Cancer |
| B1036 | 71 | 0 | 1.58 | + | + | - | 1 | DYSP |
| B1037 | 286 | 34 | 2.48 | + | + | - | 1 | DYSP |
| B1039 |  |  | 1.52 | + | + | - | 1 | DYSP |
| B1040 |  |  | 1.66 | + | + | - | 1 | DYSP |
| B1041 |  |  | 2.66 | + | + | - | 1 | DYSP |
| B1042 |  |  | 2.15 | + | + | - | 1 | DYSP |
| B1043 |  |  | 3.81 | + | + | - | 1 | DYSP |
| B1060 |  |  | 1.85 | + | + | - | 1 | DYSP |
| B1061 |  |  | 2.43 | + | + | - | 1 | DYSP |
| B1062 | 295 | 22 | 2.1 | + | + | + | 2 | Cancer |
| B1063 |  |  | 2.74 | + | - | - | 1 | GA |
| B1064 |  |  | 1.83 | + | + | - | 1 | DYSP |
| B1065 |  |  | 2.29 | + | + | - | 2 | DYSP |
| B1066 |  |  | 1.72 | + | - | - | 1 | GA |
| B1067 | 62 | 20 | 2.0 | + | + | - | 1 | DYSP |
| **Vaccine-1 (27 mice)** |  |  |  |  |  |  |  |  |
| B996 | 28 | X | 1.6 | + | - | - | 1 | GA |
| B997 |  |  | 1.5 | + | - | - | 1 | GA |
| B999 |  |  | 1.68 | + | - | - | 1 | GA |
| B1000 | 159 | 21 | 1.78 | + | + | - | 1 | DYSP |
| B1001 | 100 | 20 | 1.4 | + | - | - | 2 | DYSP |
| B1002 |  |  | 1.76 | + | - | - | 1 | GA |
| B1003 | 83 | X | 1.71 | + | - | - | 1 | GA |
| B1020 | 188 | 23 | 2.14 | + | - | - | 1 | GA |
| B1021 | 125 | X | 1.86 | + | - | - | 1 | GA |
| B1022 |  |  | 1.66 | + | - | - | 1 | GA |
| B1023 |  |  | 1.57 | + | - | - | 1 | GA |
| B1024 |  |  | 1.64 | + | - | - | 1 | GA |
| B1025 | 42 | X | 2.03 | + | - | - | 2 | GA |
| B1026 | 3050 | 31 | 2.58 | + | - | - | 1 | GA |
| B1044 |  |  | 1.73 | + | - | - | 1 | GA |
| B1045 |  |  | 1.28 | + | - | - | 1 | GA |
| B1048 | 105 | 21 | 2.39 | + | + | - | 1 | DYSP |
| B1049 |  |  | 1.91 | + | - | - | 1 | GA |
| B1050 | 175 | X | 3.55 | + | - | - | 1 | GA |
| B1051 |  |  | 2.07 | + | - | - | 1 | GA |
| B1068 | 2156 | 0 | 2.02 | + | - | - | 1 | GA |
| B1070 |  |  | 1.86 | + | + | - | 1 | DYSP |
| B1071 | 167 | 0 | 3.95 | + | - | - | 1 | GA |
| B1072 |  |  | 1.71 | + | - | - | 1 | GA |
| B1073 | 31 | X | 2.15 | + | + | - | 2 | DYSP |
| B1074 |  |  | 1.79 | + | + | - | 1 | DYSP |
| B1075 | 333 | 34 | 1.83 | + | - | - | 1 | GA |
| **Vaccine-2 (28 mice)** |  |  |  |  |  |  |  |  |
| B1004 |  |  | 1.13 | + | - | - | 1 | GA |
| B1005 |  |  | 2.39 | + | - | - | 1 | GA |
| B1006 |  |  | 1.56 | + | - | - | 1 | GA |
| B1009 |  |  | 3.66 | + | - | - | 2 | GA |
| B1010 |  |  | 1.43 | + | + | - | 2 | DYSP |
| B1011 |  |  | 1.57 | + | + | - | 1 | DYSP |
| B1028 |  |  | 1.39 | + | - | - | 1 | GA |
| B1029 |  |  | 1.61 | + | - | - | 1 | GA |
| B1030 |  |  | 1.76 | + | - | - | 1 | GA |
| B1031 |  |  | 1.76 | + | - | - | 1 | GA |
| B1032 |  |  | 2.0 | + | - | - | 1 | GA |
| B1033 |  |  | 1.38 | + | - | - | 1 | GA |
| B1034 |  |  | 1.42 | + | - | - | 1 | GA |
| B1052 |  |  | 1.47 | + | - | - | 1 | GA |
| B1054 |  |  | 1.38 | + | - | - | 1 | GA |
| B1055 | 1000 | 5 | 1.99 | + | - | - | 2 | GA |
| B1056 |  |  | 1.83 | + | - | - | 1 | GA |
| B1057 |  |  | 2.01 | + | - | - | 1 | GA |
| B1058 | 375 | 19 | 1.63 | + | - | - | 1 | GA |
| B1059 |  |  | 1.71 | + | - | - | 1 | GA |
| B1076 | 94 | X | 1.66 | + | - | - | 1 | GA |
| B1077 | 188 | 1 | 1.56 | + | - | - | 1 | GA |
| B1078 |  |  | 2.06 | + | - | - | 1 | GA |
| B1079 | 125 | 21 | 1.74 | + | - | - | 1 | GA |
| B1080 |  |  | 1.88 | + | - | - | 1 | GA |
| B1081 |  |  | 1.58 | + | - | - | 1 | GA |
| B1082 | 917 | 0 | 1.83 | + | - | - | 1 | GA |
| B1083 | 1096 | 0 | 2.0 | + | - | - | 1 | GA |
| **Non-infected Control (4 mice)** |  |  |  |  |  |  |  |  |
| B1084 |  |  | 1.38 | + | - | - |  | GA |
| B1115 |  |  | 1.49 | - | - | - |  | Healthy |
| B1116 |  |  | 4.82 | - | - | - |  | Healthy |
| B1117 |  |  | 1.46 | - | - | - |  | Healthy |

1 X – slow growing outputs

**Table S5A related to Fig. S5B.**
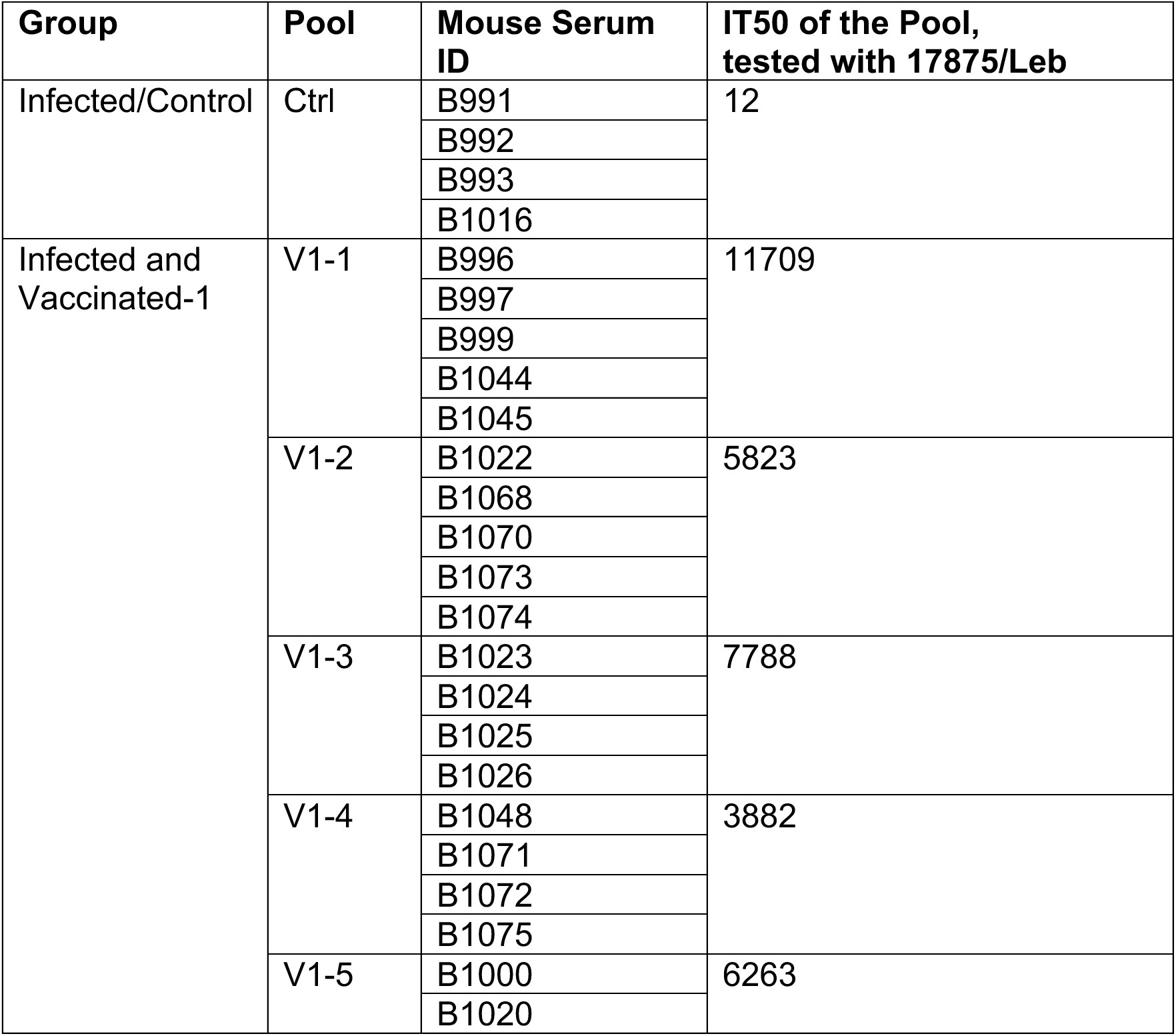
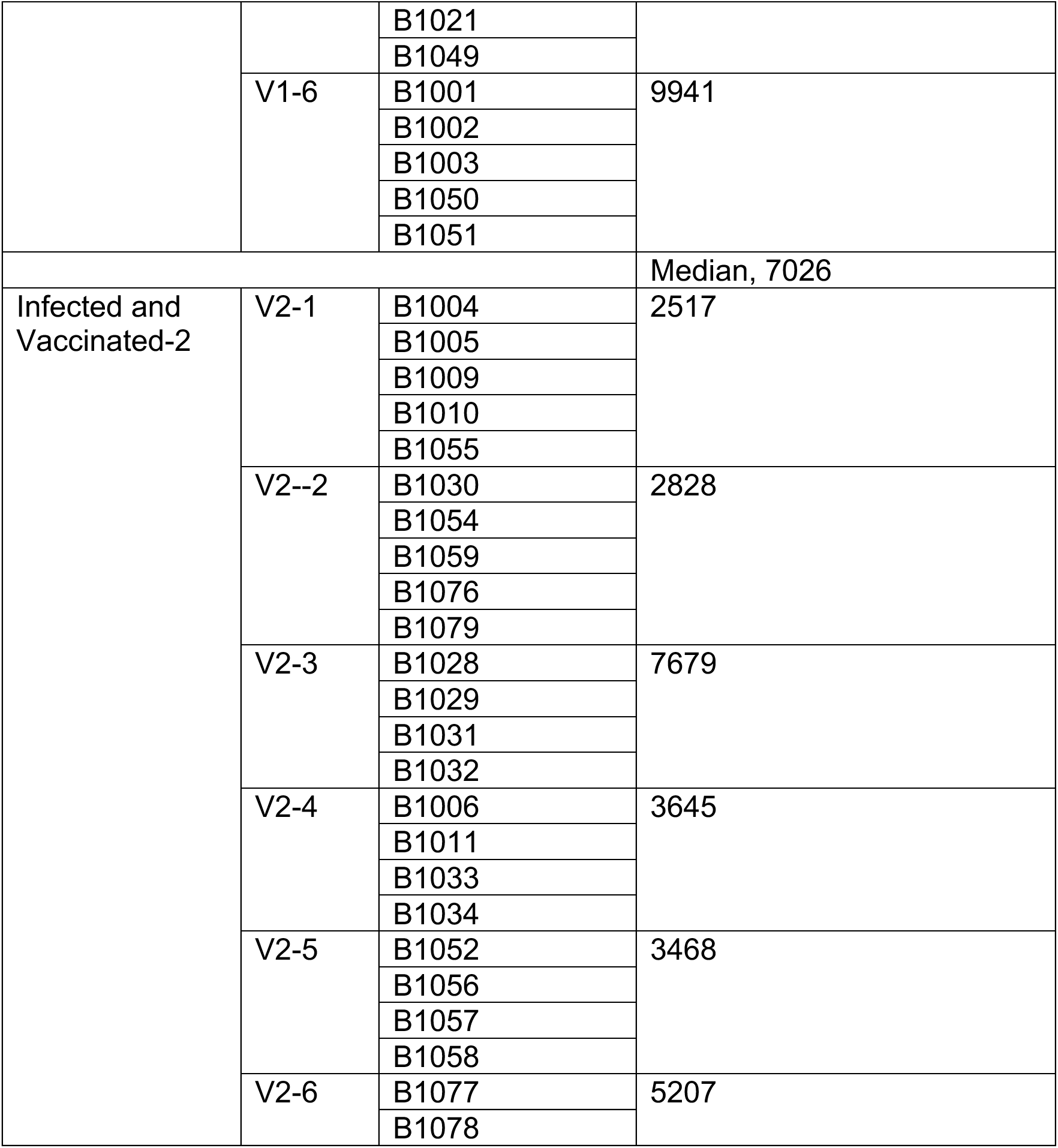
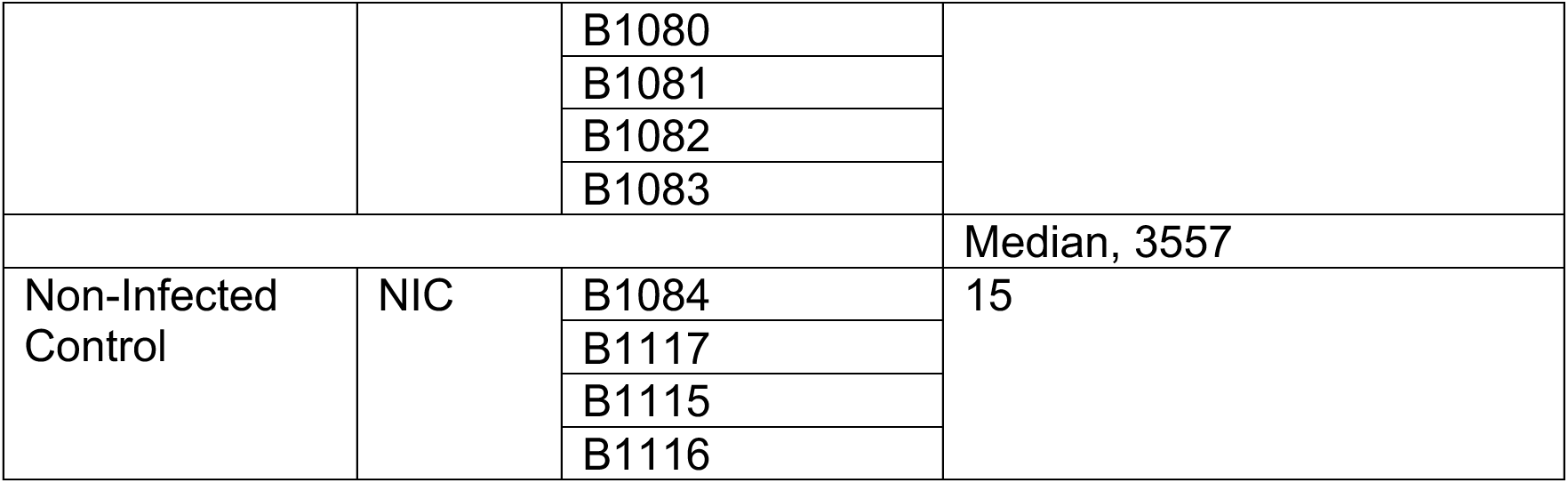
IT50 of pooled sera of infected (control) and infected and vaccinated mice, from 3^rd^ Vaccine Experiment.

**Table S5B related to Figure 6B.**
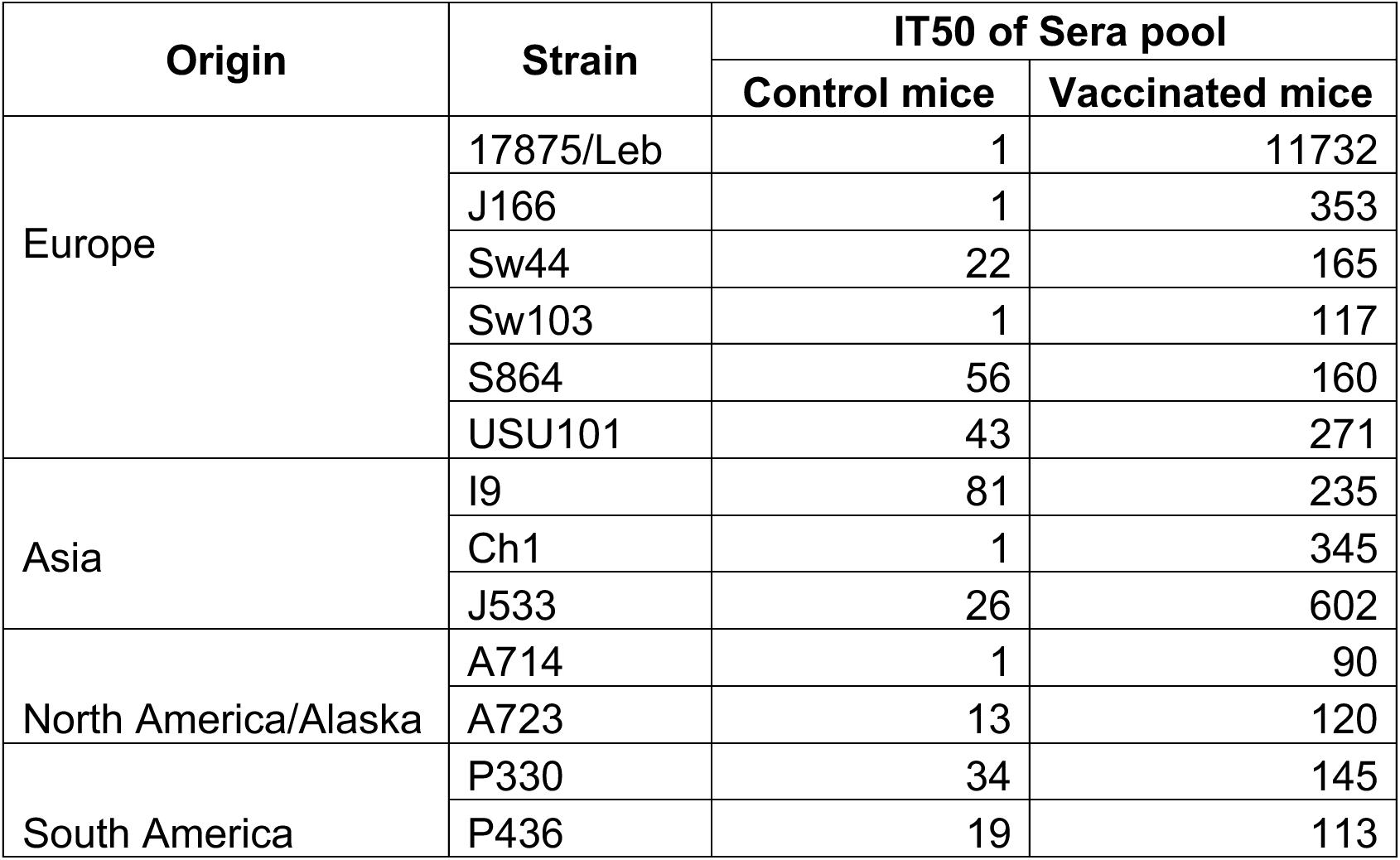
The IT50 of pooled sera from the 2^nd^ Vaccine Experiment of Infected Control mice and Vaccinated mice.

| Origin | Strain | IT50 of Sera pool |  |
| --- | --- | --- | --- |
|  |  | Control mice | Vaccinated mice |
| Europe | 17875/Leb | 1 | 11732 |
|  | J166 | 1 | 353 |
|  | Sw44 | 22 | 165 |
|  | Sw103 | 1 | 117 |
|  | S864 | 56 | 160 |
|  | USU101 | 43 | 271 |
| Asia | I9 | 81 | 235 |
|  | Ch1 | 1 | 345 |
|  | J533 | 26 | 602 |
| North America/Alaska | A714 | 1 | 90 |
|  | A723 | 13 | 120 |
| South America | P330 | 34 | 145 |
|  | P436 | 19 | 113 |

